# Cancer cells differentially modulate mitochondrial respiration to alter redox state and enable biomass synthesis in nutrient-limited environments

**DOI:** 10.1101/2025.05.09.653205

**Authors:** Sarah M. Chang, Muhammad Bin Munim, Sonia E. Trojan, Huel Cox, Anna Shevzov-Zebrun, Keene L. Abbott, Renee Chang, Matthew G. Vander Heiden

## Abstract

The cell NAD+/NADH ratio can constrain biomass synthesis and influence proliferation in nutrient-limited environments. However, which cell processes regulate the NAD+/NADH ratio is not known. Here, we find that some cancer cells elevate the NAD+/NADH ratio in response to serine deprivation by increasing mitochondrial respiration. Cancer cells that elevate mitochondrial respiration have higher serine production and proliferation in serine limiting conditions than cells with no mitochondrial respiration response, independent of serine synthesis enzyme expression. Increases in mitochondrial respiration and the NAD+/NADH ratio promote serine synthesis regardless of whether serine is environmentally limiting. Lipid deprivation can increase the NAD+/NADH ratio via mitochondrial respiration in some cells, including cells that do not increase respiration following serine deprivation. Thus, in cancer cells where lipid depletion raises the NAD+/NADH ratio, proliferation in serine depleted environments improves when lipids are also depleted. Taken together, these data suggest that changes in mitochondrial respiration in response to nutrient deprivation can influence the NAD+/NADH ratio in a cell-specific manner to impact oxidative biomass synthesis and proliferation. Given the complexity of tumor microenvironments, this work provides a metabolic framework for understanding how levels of more than one environmental nutrient affect cancer cell proliferation.

## Introduction

Rapidly proliferating cells, including cancer cells, must acquire biomass precursors such as amino acids, lipids, and nucleotides to support cell growth and division (Deberardinis, 2008; Faubert, 2020; Hanahan, 2011; Pavlova, 2017; Vander Heiden, 2017). While cells can obtain biomass precursors from the environment, many tissue and tumor environments can be deficient in necessary nutrients (Cognet, 2024; Gullino, 1964; Lyssiotis, 2017). In these conditions, cells depend on synthesis pathways to accumulate sufficient biomass for proliferation (Apiz Saab, 2023; Ferraro, 2021; Ngo, 2020; Sciacovelli, 2022; Sullivan, 2019b; Sullivan, 2019a). Synthesis of many biomass precursors involve oxidation reactions that require the redox cofactor NAD+ to act as an electron acceptor. There is accumulating evidence that NAD+ availability and the cell redox state (NAD+/NADH ratio) can limit the synthesis of biomass precursors, including aspartate (Birsoy, 2015; Sullivan, 2015), asparagine (Krall, 2021), fatty acids (Li, 2022), serine (Baksh, 2020; Bao, 2016; Broeks, 2023; Diehl, 2019), and nucleotides (Bao, 2016; Diehl, 2019; Wu, 2024). Thus, the NAD+/NADH ratio impacts cancer cell proliferation in nutrient environments that increase dependence on biosynthetic oxidation reactions. However, what determines the cell NAD+/NADH ratio and whether the NAD+/NADH ratio is modulated in response to environmental nutrient availability is not well characterized. Moreover, whether differences in the NAD+/NADH ratio across cell types affect sensitivity to different nutrient limitations is not known.

Some tumor microenvironments have low levels of serine, an amino acid used to produce many biomass components, including proteins, nucleotides, glutathione, and lipids. Thus, low serine environments can limit cancer cell proliferation and tumor growth (Banh, 2020; Maddocks, 2013; 2017; Ngo, 2020; Sullivan, 2019b). In serine depleted environments, cancer cells must increase serine synthesis to maintain proliferation (DeNicola, 2015; Labuschagne, 2014; Locasale, 2011; Possemato, 2011; Sullivan, 2019b). Serine synthesis involves the conversion of the glycolytic intermediate 3-phosphoglycerate (3-PG) into serine. The production of 3-PG from glucose involves an oxidation reaction catalyzed by glyceraldehyde 3-phosphate dehydrogenase (GADPH). 3-PG is then oxidized by phosphoglycerate dehydrogenase (PHGDH) in the first step of the serine synthesis pathway. Both GADPH and PHGDH require NAD+ as an electron acceptor to enable serine synthesis. Experimentally blunting NAD+ regeneration from NADH reduces serine synthesis and is rescued by oxidizing the NAD+/NADH ratio via exogenous electron acceptor supplementation (Bao, 2016; Diehl, 2019; Gravel, 2014). Similarly, de novo fatty acid synthesis can be constrained by the NAD+/NADH ratio, as the production of fatty acids involves multiple oxidation reactions to generate citrate from either glucose or glutamine. Thus, decreasing the NAD+/NADH ratio sensitizes cells to environmental lipid depletion (Li, 2022). Together, these findings demonstrate that exogenous changes to the NAD+/NADH ratio can alter serine and citrate production and impact proliferation in serine- or lipid-depleted environments. However, auxotrophy for a single nutrient is not sufficient to determine whether cancer cells can form tumors in a tissue, where levels of variable nutrients may be environmentally limiting for proliferation (Abbott, 2026; Sivanand, 2024; Sullivan, 2019a).Thus, we aimed to uncover how the endogenous NAD+/NADH ratio of cancer cells is regulated to influence biosynthesis in different nutrient depleted environments and its role in determining cancer cell proliferation when multiple oxidized nutrients are limiting.

By culturing cancer cells in serine depleted conditions and examining the relationship between the NAD+/NADH ratio and serine synthesis, we find that the NAD+/NADH ratio is elevated in response to serine deprivation in some but not all cancer cells. The elevated NAD+/NADH ratio is driven by increased mitochondrial respiration and promotes serine synthesis to confer resistance to environmental serine depletion. A similar relationship between mitochondrial respiration and the NAD+/NADH ratio was also observed in lipid depleted environments, where higher mitochondrial respiration in some cells led to elevated NAD+/NADH ratios and enhanced citrate synthesis. Lastly, we find that any perturbation that increases the NAD+/NADH ratio, including lipid deprivation, could paradoxically improve the proliferation of cells in serine depleted conditions. Together, these data reveal that the NAD+/NADH ratio is regulated by mitochondrial respiration in response to environmental nutrients in a cancer cell-specific manner and that this influences oxidative biosynthesis and proliferation. These data also provide insight into how levels of multiple nutrients in tissue microenvironments may cooperate to enable biomass synthesis and cell proliferation.

## Results

### Modulating the cell NAD+/NADH ratio proportionally alters serine synthesis

To begin investigating the processes that determine the cell NAD+/NADH ratio and influence biomass synthesis, we examined the relationship between the NAD+/NADH ratio and serine synthesis in serine-replete conditions (400 μM serine, the concentration of serine in DMEM) and serine depleted conditions (DMEM media formulated without serine). Previous studies have found that decreasing the cell NAD+/NADH ratio (more reduced) can suppress serine production (Baksh, 2020; Bao, 2016; Broeks, 2023; Diehl, 2019). To confirm this, we varied the cell NAD+/NADH ratio upon serine withdrawal and measured the serine synthesis rate of A549 non-small cell lung cancer cells, which transcriptionally upregulate the serine synthesis enzyme PHGDH (DeNicola, 2015). We increased the NAD+/NADH ratio by treating cells with the exogenous electron acceptor alpha-ketobutyrate (AKB), which can be reduced to alpha-hydroxybutyrate to regenerate NAD+ (Sullivan, 2015) (**Figure 1A**). We decreased the NAD+/NADH ratio by treating cells with rotenone, an inhibitor of mitochondrial complex I, which regenerates NAD+ in support of respiration. AKB and rotenone dose-dependently increased and decreased the cell NAD+/NADH ratio, respectively (**Figure 1B**). To measure the serine synthesis rate in cells, we performed kinetic isotope tracing using uniformly ^13^C-labeled glucose (U-^13^C-glucose) and measured production of M+3 labeled serine (**Figure 1C**). Because intracellular serine, including newly synthesized serine, rapidly exchanges with the extracellular environment (Labuschagne, 2014), we collected both media and cells to help improve the detection of newly synthesized serine produced from glucose. We find that increasing the cell NAD+/NADH ratio with AKB led to proportionally higher serine synthesis rates whereas lowering the NAD+/NADH ratio with rotenone led to proportionally lower serine synthesis rates (**Figure 1D**). This was not explained by changes in PHGDH protein expression, as altering the NAD+/NADH ratio with AKB or rotenone did not affect PHGDH protein levels despite modulating serine synthesis (**Supplementary Figure 1A,B**). Additionally, neither rotenone treatment nor serine deprivation had a large effect on cell viability (**Supplementary Figure 1C**). Of note, we find that serine synthesis rate is positively correlated with the cell NAD+/NADH ratio (**Figure 1E**). Together, these data demonstrate that modulating the NAD+/NADH ratio leads to proportional changes in serine synthesis rates.

**Figure 1.**
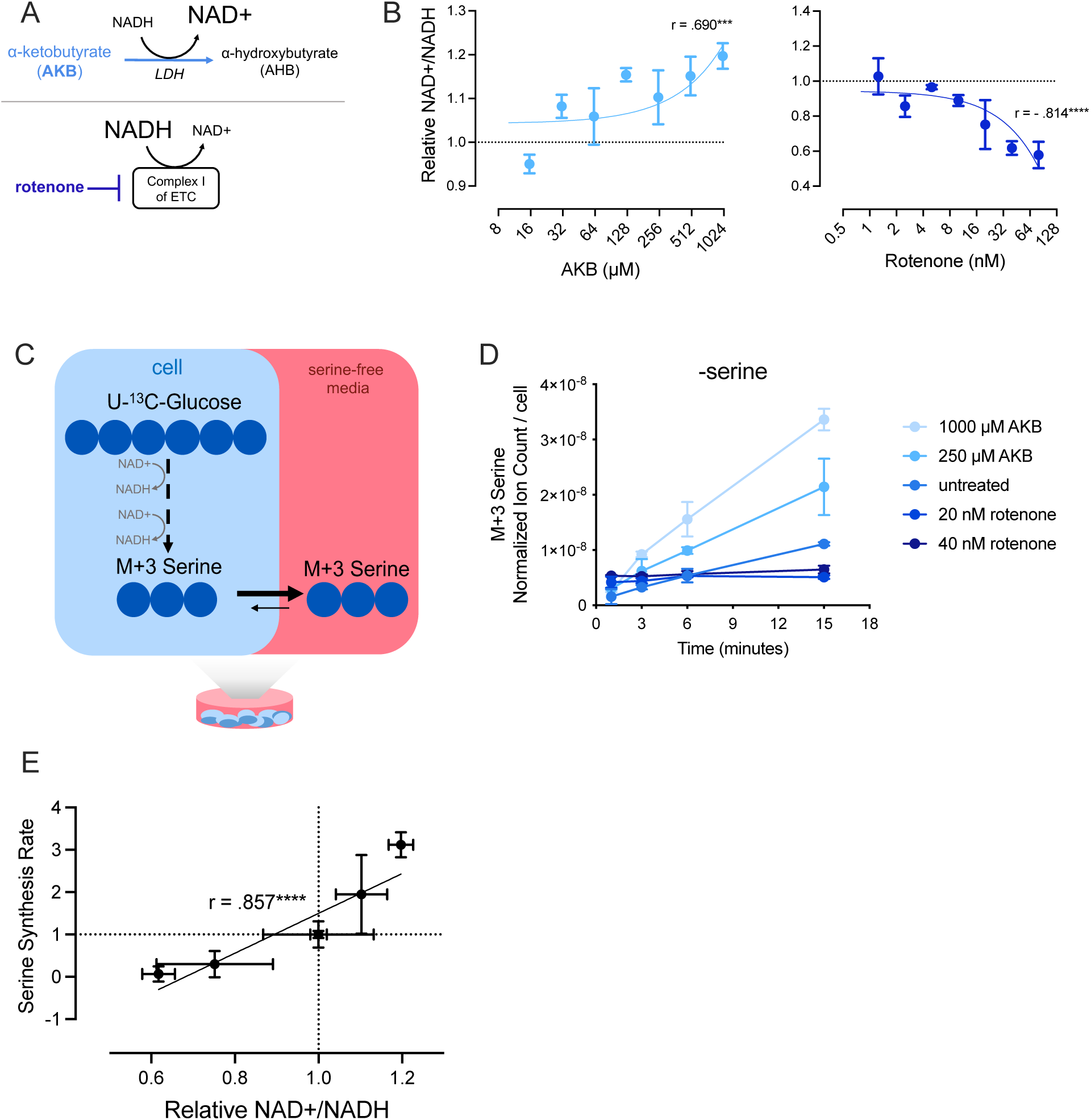
The NAD+/NADH ratio is proportional to serine synthesis rate. **(A)** α-ketobutyrate (AKB) is an exogenous electron acceptor that promotes the oxidation of NADH into NAD+ through its conversion to α-hydroxybutyrate (AHB). Rotenone inhibits activity of complex I of the mitochondrial electron transport chain (ETC) to block NADH oxidation into NAD+. **(B)** NAD+/NADH ratio measured in A549 cells cultured without serine and indicated concentrations of AKB (left) and rotenone (right) for 24 hours. NAD+/NADH ratios are normalized to untreated cells cultured without serine, n=3. Pearson correlation coefficients and P-values were calculated by simple linear regression, ***p<0.005, ****p<0.001. **(C)** Schematic depicting NAD+ requirement to produce serine labeled (M+3) from U-^13^C-glucose. Under culture conditions without serine, intracellular serine rapidly effluxes from cells into culture media. Thus, cells and media were jointly analyzed at each time point of kinetic tracing to better capture levels of newly synthesized serine. **(D)** Serine labeled (M+3) from U-^13^C-glucose over time in A549 cells cultured without serine and with indicated concentrations of AKB or rotenone for 24 hours prior to U-^13^C-glucose exposure. Serine levels are normalized to internal norvaline standard and cell number measured in indicated conditions, n=3. **(E)** Relative NAD+/NADH ratios from A549 cells cultured without serine after exposure to 250 μΜ AKB, 1000 μΜ AKB, 20 nM rotenone, or 40 nM rotenone for 24 hours are plotted against the serine synthesis rates from corresponding conditions as shown in (D). Serine synthesis rates are calculated from labeled serine (M+3) after 1 and 15 minutes of U-^13^C-glucose exposure and normalized to untreated culture conditions without serine, n=3. Pearson correlation coefficients and P-values were calculated by simple linear regression, ****p<0.001. Data shown for all panels represent mean ± SD.

### Cancer cells with elevated NAD+/NADH ratios following serine withdrawal exhibit greater serine synthesis

Based on our finding that the NAD+/NADH ratio correlates with serine synthesis rate, we wondered whether endogenous cell NAD+/NADH ratios, which can differ across cancer cells, are predictive of proliferation rate following serine deprivation. To test this, we measured how proliferation rate varied in a panel of cancer cells derived from various tissues-of-origin and with different genetic driver mutations, and found a wide range of sensitivity to serine deprivation (**Figure 2A**, **Supplementary Figure 2A, Supplementary Table 1**). Because PHGDH expression is important for proliferation in serine depleted conditions (DeNicola, 2015; Locasale, 2011; Maddocks, 2017; Possemato, 2011; Sullivan, 2019b), we first assessed the protein expression of the serine synthesis enzymes PHGDH, PSAT1, and PSPH across the different cancer cells to examine how well this explained environmental serine dependence. As expected, cells with low serine synthesis enzyme protein levels (MCF7, MDA-MB-231) were most sensitive to serine withdrawal while cells with higher serine synthesis enzyme protein levels were more resistant to serine withdrawal (PHGDH-overexpressing MDA-MB-231, A375) (**Supplementary Figure 2A-C**). However, for many cancer cells (Calu6, A549, 8988T, MIA PaCa-2, H1299, HCT116), serine synthesis enzyme expression alone could not fully explain sensitivity to serine withdrawal (**Supplementary Figure 2C**). For example, when comparing the non-small cell lung cancer cells A549 and H1299, H1299 cells express less PHGDH protein and similar PSAT1 and PSPH protein levels than A549 cells but are more resistant to serine deprivation **(Supplementary Figure 2B-E)**. Consistent with lower PHGDH expression, H1299 cells express lower levels of NRF2, a transcriptional regulator of serine biosynthesis enzymes (DeNicola, 2015). When examining the cancer cells where serine synthesis enzyme expression could not fully explain the differences in proliferation upon serine depletion, there was a surprising negative correlation between proliferation and PHGDH protein expression, no relationship between proliferation and PSAT1 protein expression, and a positive correlation between proliferation and PSPH protein expression (**Supplementary Figure 2F**). Given our finding that the NAD+/NADH ratio correlates with serine synthesis rate, we hypothesized that differences in the NAD+/NADH ratio upon serine deprivation could contribute to the variability in sensitivity to serine withdrawal across cells with similar serine synthesis pathway protein levels. To test this, we measured the cell NAD+/NADH ratios in the presence and absence of exogenous serine. Interestingly, we find that serine starvation elevated the NAD+/NADH ratio in some cells, which we term “redox responder cells” and label in blue. In contrast, we also find that certain cancer cells minimally change the NAD+/NADH ratio in response to serine depletion, which we term “redox non-responder cells” and label in yellow (**Figure 2B, Supplementary Figure 2G**). Of note, whether the NAD+/NADH ratio of a cell was more or less oxidized in serine-replete conditions was not predictive of response to serine withdrawal (**Supplementary Figure 2G)**. Strikingly, redox responder cells were more resistant to serine depletion compared to redox non-responder cells (**Figure 2C**). The NAD+/NADH ratio was also elevated in cancer cells with higher serine synthesis enzyme levels upon serine depletion and were the most resistant to serine depletion (A375, PHGDH-overexpressing MDA-MB-231) (**Supplementary Figure 2B,C,G**). To confirm that an elevated NAD+/NADH ratio in response to serine depletion was associated with elevated serine synthesis, we compared how serine synthesis rates changed following serine withdrawal in A549 and H1299 cells. We performed focused comparisons between A549 and H1299 cells because they exhibit differences in proliferation upon serine deprivation that are not explained by PHGDH protein expression, demonstrate differing responses of the cell NAD+/NADH ratio upon serine deprivation, and have similar basal proliferation rates. We find that after 24 hours in serine depleted conditions, H1299 cells demonstrated a higher rate of serine synthesis, a greater increase in serine synthesis, and greater intracellular serine levels than A549 cells, consistent with the elevated NAD+/NADH ratio in H1299 cells (**Figure 2D,E; Supplementary Figure 2H,I**). Importantly, we confirmed kinetic U-^13^C-glucose tracing was performed at metabolic steady state by ensuring metabolite levels were stable at each collected time point (**Supplementary Figure 2I**). Taken together, these data suggest that in some cells, the cell NAD+/NADH ratio is responsive to serine availability. Moreover, increases in the NAD+/NADH ratio correlate with increased serine synthesis and proliferation rates in serine depleted conditions independent of PHGDH protein expression.

**Figure 2.**
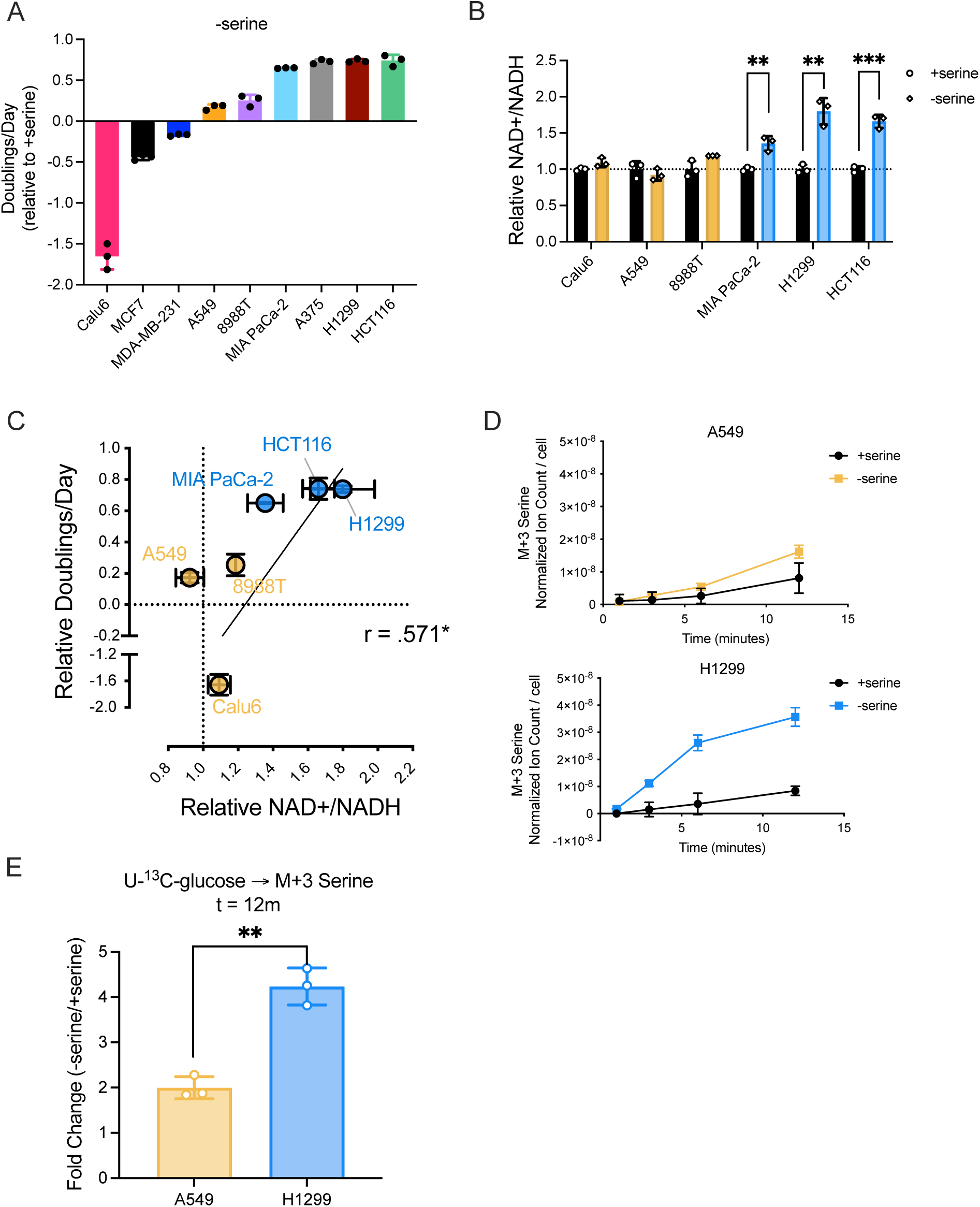
The NAD+/NADH ratio differs between cancer cells upon serine withdrawal and correlates with proliferation in serine depleted conditions. **(A)** Proliferation rate (doublings per day) of indicated cancer cells cultured without serine for 72 hours normalized to proliferation rate of corresponding cancer cells cultured with serine, n=3. **(B)** NAD+/NADH ratios of indicated cells cultured with or without serine for 24 hours. Yellow indicates cells with unaltered NAD+/NADH ratios (redox non-responder cells) and blue indicates cells with elevated NAD+/NADH ratios when cultured without serine (redox responder cells). Values are normalized to the NAD+/NADH ratios of cells cultured with serine, n=3. P-values were calculated by unpaired Student’s t-test, **p<0.01, ***p<0.005. **(C)** Relative NAD+/NADH ratios of cells cultured without serine are plotted against proliferation rates of corresponding cells cultured without serine. Data points are labeled to denote the different cell lines (Calu6, A549, 8988T, MIA PaCa-2, H1299, and HCT116). Pearson correlation coefficient and P-values were calculated by simple linear regression, *p<0.05. **(D)** Serine labeled (M+3) from U-^13^C-glucose over time in A549 and H1299 cells cultured with or without serine for 24 hours prior to U-^13^C-glucose exposure. Serine levels were normalized to internal norvaline standard and cell number measured in indicated conditions, n=3. **(E)** Absolute fold change of serine labeled (M+3) from U-^13^C-glucose in A549 and H1299 cells cultured without serine relative to serine labeled (M+3) from U-^13^C-glucose in A549 and H1299 cells cultured with serine for 24 hours after 12 minutes of U-^13^C-glucose exposure, n=3. P-values were calculated by unpaired Student’s t-test, **p<0.01. Data shown for all panels are means ± SD.

### Increased mitochondrial respiration is associated with an elevated NAD+/NADH ratio and increased serine synthesis

The finding that some cancer cells exhibit a more oxidized NAD+/NADH ratio upon serine starvation is surprising, particularly when increased serine synthesis will result in a higher demand for NAD+ to support increased flux through GAPDH and PHGDH (Diehl, 2019; Maddocks, 2017). We hypothesized that a more oxidized NAD+/NADH ratio could support greater serine synthesis and thus sought to identify the processes that increase the NAD+/NADH ratio in some but not all cancer cells. Lactate production via lactate dehydrogenase (LDH) is a major NAD+ regenerating process, particularly in cancer cells with high glucose fermentation (Fantin, 2006; Luengo, 2021; Wang, 2022) (**Supplementary Figure 3A**); however, lactate secretion relative to glucose consumption was not elevated following serine deprivation in any cells tested (**Supplementary Figure 3B**). The NAD+ salvage pathway regenerates NAD+ from nicotinamide via nicotinamide phosphoribosyl transferase (NAMPT) and nicotinamide mononucleotide adenylyltransferase (NMNAT) and can be important for supporting PHGDH-driven serine synthesis (**Supplementary Figure 3A**) (Murphy, 2018). While A549 cells were more sensitive to the NAMPT inhibitor FK866 in serine-replete conditions, there was no difference in the sensitivity of A549 and H1299 cells to FK866 in serine depleted conditions (**Supplementary Figure 3C**). These data suggest that redox responder and non-responder cells do not have differential dependency on the NAD+ salvage pathway when cultured without serine. Moreover, supplementing redox non-responder A549 cells with nicotinamide mononucleotide (NMN), a precursor to NAD+, did not improve proliferation in serine depleted conditions (**Supplementary Figure 3D**). Mitochondrial respiration is another major way cells regenerate NAD+ from NADH (Handy, 2012). Thus, we tested whether changes in mitochondrial respiration, as measured by cellular oxygen consumption rate (OCR), accompany the increase in the NAD+/NADH ratio following serine withdrawal. Indeed, cells that increased their NAD+/NADH ratio following serine deprivation also exhibited increased OCR, whereas cells with unaltered NAD+/NADH ratios following serine withdrawal did not (**Figure 3A, Supplementary Figure 3E**). To test whether elevated oxygen consumption was related to serine deprivation, we cultured H1299 and A549 cells in varying amounts of extracellular serine and measured OCR. Oxygen consumption of A549 cells was not impacted by varying extracellular serine levels, while oxygen consumption of H1299 cells increased as extracellular serine decreased below 100 μM, the serine concentration where proliferation begins to be affected by serine limitation (**Figure 3B, Supplementary Figure 3F**). The increase in OCR also corresponded with the NAD+/NADH ratio at different serine levels in H1299 cells, while the NAD+/NADH ratio in A549 cells was unchanged by extracellular serine availability (**Figure 3C, D).** Consistently, the NAD+/NADH ratio of H1299 cells cultured in different concentrations of serine positively correlated with serine synthesis, whereas A549 cells displayed minimal changes in serine synthesis rate (**Figure 3E, F**). Notably, PHGDH protein levels remained constant as extracellular serine levels were decreased in both A549 and H1299 cells (**Supplementary Figure 3G**). Together, these findings suggest that elevated NAD+/NADH ratios in response to decreasing environmental serine are linked to elevated mitochondrial respiration in select cancer cells and correlate with increased serine synthesis.

**Figure 3.**
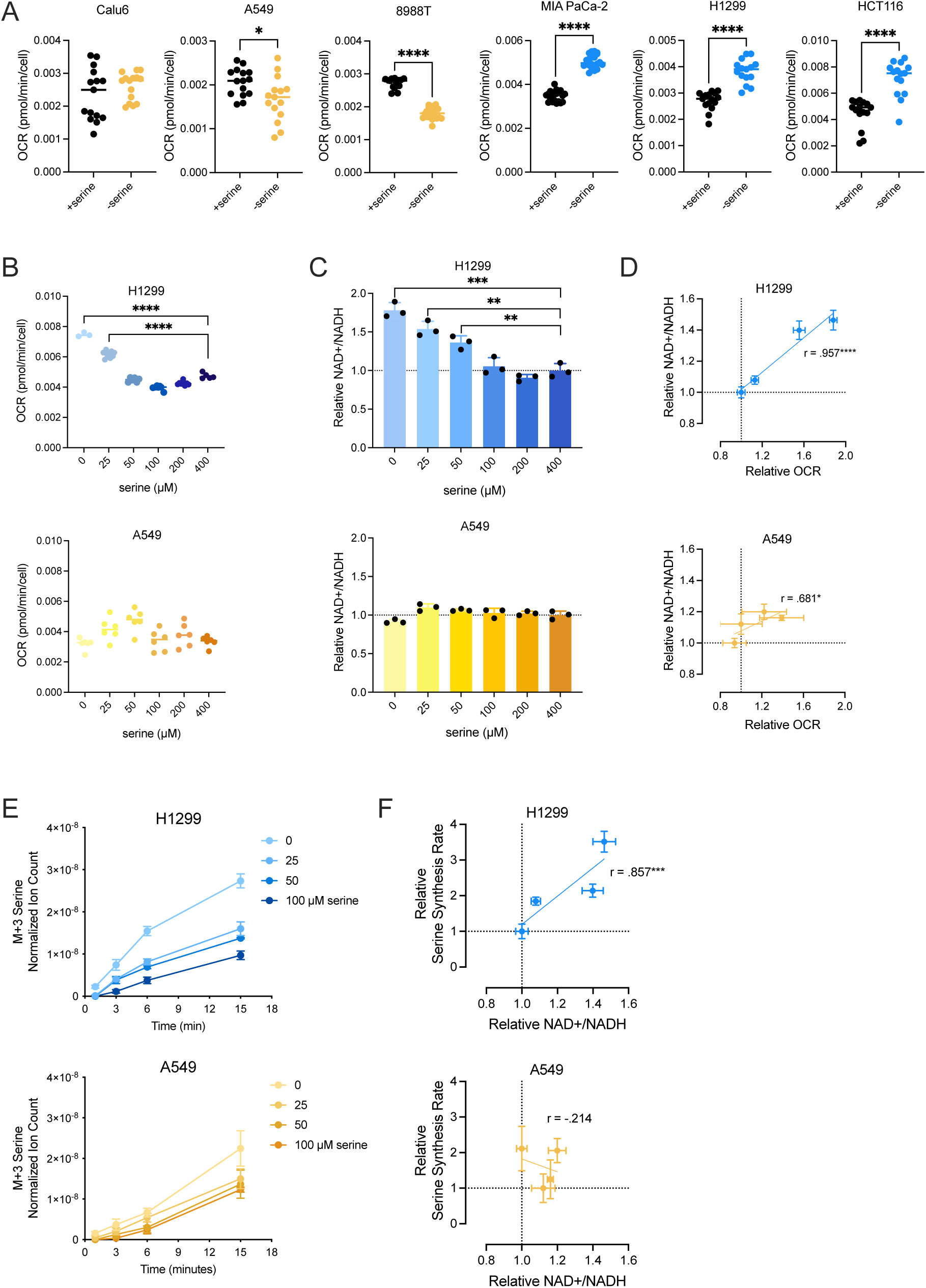
Elevated mitochondrial respiration is associated with increased NAD+/NADH ratios and serine synthesis rates. **(A)** Oxygen consumption rate (OCR) of indicated cells cultured with or without serine for 24 hours. Values are averages of three repeat measurements, n=14-20. P-values were calculated by unpaired Student’s t-test, *p<0.05, ****p<0.001. **(B)** OCR of cells (H1299 top, A549 bottom) cultured with indicated concentrations of serine for 24 hours. Values are averages of three repeat measurements, n=3-9. P-values were calculated by unpaired Student’s t-test, ****p<0.001. **(C)** NAD+/NADH ratios of cells (H1299 top, A549 bottom) cultured with indicated concentrations of serine for 24 hours. Values are normalized to the NAD+/NADH ratio of cells cultured in 400 μM serine (serine concentration in DMEM), n=3. P-values were calculated by unpaired Student’s t-test, **p<0.01, ***p<0.005. **(D)** Relative OCR plotted against relative NAD+/NADH ratios in H1299 and A549 cells cultured in 0, 25, 50, and 100 μM serine for 24 hours. OCR and NAD+/NADH ratios are normalized to measurements from the 100 μΜ serine condition, n=3. Pearson correlation coefficients and P-values were calculated by simple linear regression, *p<0.05, ****p<0.001. **(E)** Serine labeled (M+3) from U-^13^C-glucose over time in cells (H1299 top, A549 bottom) cultured with indicated serine concentrations for 24 hours prior to U-^13^C-glucose exposure. Serine levels are normalized to internal norvaline standard and cell number measured in indicated conditions, n=3. **(F)** Relative NAD+/NADH ratios of cells (H1299 top, A549 bottom) cultured with 0, 25, 50, and 100 μM serine for 24 hours plotted against relative serine synthesis rates of cells cultured in corresponding conditions. Serine synthesis rates are calculated from labeled serine (M+3) after 1 and 15 minutes of U-^13^C-glucose exposure and normalized to untreated culture conditions without serine, n=3. Pearson correlation coefficients and P-values were calculated by simple linear regression, ***p<0.005. Data shown for all panels are means ± SD.

### Mitochondrial respiration governs the cell NAD+/NADH ratio and influences serine synthesis rate

To test whether increased mitochondrial respiration following serine depletion causes an increase in the cell NAD+/NADH ratio, we treated redox non-responder cells with the proton ionophore FCCP (trifluoromethoxy carbonylcyanide phenylhydrazone), which pharmacologically increases mitochondrial oxygen consumption by uncoupling respiration from ATP synthesis (McLaughlin, 1980; Terada, 1990). Increasing respiration with FCCP elevates mitochondrial NAD+ regeneration and the cell NAD+/NADH ratio (Luengo, 2021) (**Figure 4A**). Indeed, FCCP treatment improved proliferation of redox non-responder cells in serine depleted conditions (**Figure 4B, Supplementary Figure 4A**). In contrast, FCCP did not improve the proliferation of redox responder cells cultured without serine (**Figure 4C, Supplementary Figure 4B**). Consistent with its impact on proliferation, FCCP treatment led to higher serine synthesis rates only in redox non-responder A549 cells while having no impact on the serine synthesis rates of redox responder H1299 cells (**Figure 4D,E, Supplementary Figure 5A-D**). As expected, uncoupling mitochondrial respiration from ATP synthase with FCCP led to elevated oxygen consumption in serine depleted A549 cells (**Figure 4F).** FCCP also increased oxygen consumption in serine depleted H1299 cells (**Supplementary Figure 4C**). Interestingly, FCCP raised the cell NAD+/NADH ratios of both A549 and H1299 cells following serine withdrawal despite only improving serine synthesis and proliferation in A549 cells (**Figure 4G, Supplementary Figure 4D**). The lack of impact on serine synthesis by FCCP in H1299 cells suggests that NAD+ availability may not further constrain serine synthesis in cells that already elevate mitochondrial respiration following environmental serine depletion. In this context, we considered whether another constraint such as PHGDH protein expression was limiting serine synthesis in redox responder cells treated with FCCP. Indeed, overexpressing PHGDH in serine depleted redox responder H1299 and HCT116 cells fully restored proliferation to that observed in serine-replete conditions (**Supplementary Figure 4E,G**). In contrast, proliferation of the redox non-responder cells A549 and Calu6 following serine withdrawal had the greatest improvement with simultaneous PHGDH overexpression and FCCP treatment (**Supplementary Figure 4F,G**).

**Figure 4.**
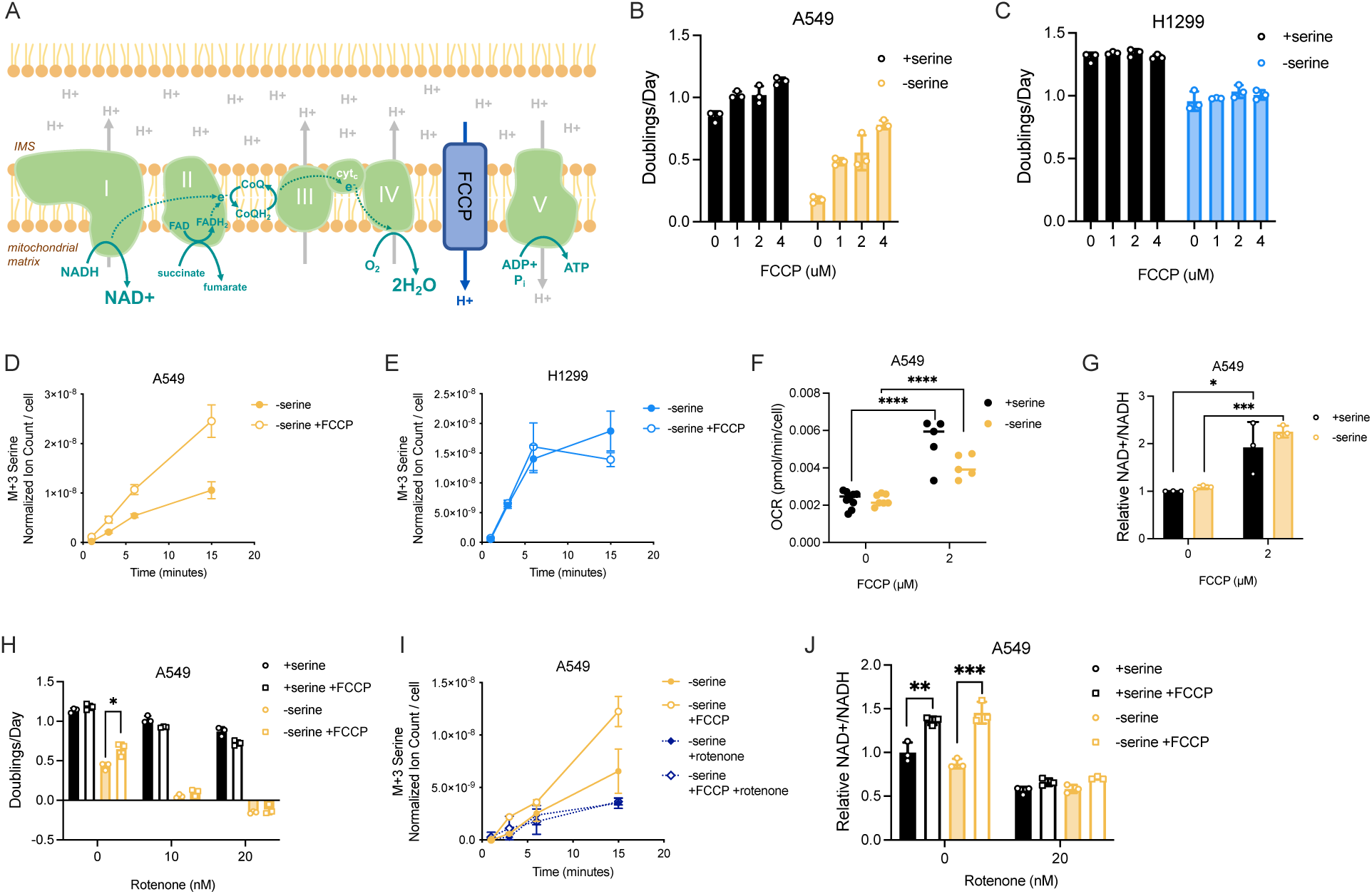
Mitochondrial respiration governs the endogenous cell NAD+/NADH ratio and influences serine synthesis. **(A)** Schematic depicting the mitochondrial electron transport chain (ETC) and the protonophore activity of FCCP that uncouples the ETC from ATP synthase activity. CoQ(H_2_) indicates the oxidized and reduced (H_2_) forms of coenzyme Q, cyt_c_ denotes cytochrome c. **(B,C)** Proliferation rate (doublings per day) of A549 (B) and H1299 cells (C) cultured with or without serine with indicated concentrations of FCCP for 72 hours, n=3. Increasing doses of FCCP statistically increased proliferation of A549 cells cultured without serine (p<0.001). P-values were calculated by simple linear regression. **(D)** Serine labeled (M+3) from U-^13^C-glucose over time in A549 cells without serine and with or without 2 μM FCCP treatment for 24 hours prior to U-^13^C-glucose exposure. Serine levels are normalized to internal norvaline standard and cell number measured in indicated conditions, n=3. **(E)** Serine labeled (M+3) from U-^13^C-glucose over time in H1299 cells without serine, and with or without 2 μM FCCP treatment for 24 hours prior to U-^13^C-glucose exposure. Serine levels are normalized to internal norvaline standard and cell number measured in indicated conditions, n=3. **(F)** Oxygen consumption rate (OCR) of A549 cells cultured with or without serine with indicated FCCP treatment for 24 hours. Values are averages of three repeat measurements, n=5-9. P-values were calculated by unpaired Student’s t-test, ****p<0.001. **(G)** Relative NAD+/NADH ratio of A549 cells cultured with or without serine with indicated FCCP treatment for 24 hours. NAD+/NADH ratios are normalized to the NAD+/NADH ratio in A549 cells cultured with serine, n=3. P-values were calculated by unpaired Student’s t-test, *p<0.05, ***p<0.005. **(H)** Proliferation rate (doublings per day) of A549 cells cultured with or without serine, 2 μM FCCP, and indicated concentrations of rotenone for 72 hours, n=3. P-values were calculated by unpaired Student’s t-test, *p<0.05. **(I)** Serine labeled (M+3) from U-^13^C-glucose over time in A549 cells cultured without serine and with or without 2 μM FCCP or 20 nM rotenone as indicated for 24 hours prior to U-^13^C-glucose exposure. Serine levels are normalized to internal norvaline standard and cell number measured in indicated conditions, n=3. **(J)** Relative NAD+/NADH ratio of A549 cells cultured with or without serine, 2 μM FCCP, or 20 nM rotenone as indicated. NAD+/NADH ratios are normalized to the NAD+/NADH ratio in A549 cells cultured with serine, n=3. P-values were calculated by unpaired Student’s t-test, **p<0.01, ***p<0.005. Data shown for all panels are means ± SD.

This underscores that changes in both the cell NAD+/NADH ratio and relevant enzyme expression impact serine synthesis in serine depleted environments. As an orthogonal approach to FCCP, we constitutively expressed the NADH oxidase from *Lactobacillus brevis* (*Lb*NOX) in both the cytoplasm and the mitochondria of A549 cells (**Supplementary Figure 6A**). We reasoned that if expressing *Lb*NOX in either the cytoplasm or the mitochondrial were to raise the cell NAD+/NADH ratio, proliferation of A549 cells upon serine depletion would be improved. To ensure *Lb*NOX activity was present in cells, we treated cells with rotenone and measured proliferation. As expected, we observed a proliferation improvement when *Lb*NOX was expressed in either the cytoplasm or the mitochondria (**Supplementary Figure 6B**). However, A549 cyto*Lb*NOX and mito*Lb*NOX cells did not exhibit an elevated cell NAD+/NADH ratio in either serine-replete or serine depleted conditions (**Supplementary Figure 6C**), consistent with prior observations (Titov, 2016). There was also no change in proliferation upon serine withdrawal when *Lb*NOX was expressed in the cytoplasm or mitochondria (**Supplementary Figure 6D**).

Because FCCP treatment led to a robust improvement in proliferation of redox non-responder cells cultured without serine, we aimed to test whether FCCP improved serine synthesis and proliferation of redox non-responder cells through oxidizing the NAD+/NADH ratio. We treated cells with the mitochondrial complex I inhibitor rotenone in conjunction with FCCP. Rotenone treatment ablated the ability of FCCP to improve both proliferation and serine synthesis and blocked the ability of FCCP to increase the NAD+/NADH ratio (**Figure 4H-J, Supplementary Figure 5E,F**). In contrast, treating cells with oligomycin, an ATP synthase inhibitor, did not affect the ability of FCCP to improve proliferation (**Supplemental Figure 4H,I**). This supports the conclusion that FCCP increases serine synthesis and proliferation in serine depleted environments by increasing the NAD+/NADH ratio via mitochondrial complex I-mediated NAD+ regeneration. Moreover, FCCP treatment neither increased PHGDH protein expression nor glucose consumption, indicating that improved proliferation and serine synthesis is not caused by changes in enzyme expression or carbon availability for serine synthesis (**Supplementary Figure 4J,K**). Together, these data argue that mitochondrial respiration, either endogenously increased in response to serine deprivation, or as a result of FCCP treatment, can govern the cell NAD+/NADH ratio and directly influence the serine synthesis rate in response to serine withdrawal.

### The malate aspartate shuttle enzymes MDH1 and GOT2 support elevated mitochondrial respiration in redox responder H1299 cells in serine-depleted environments

The malate aspartate shuttle (MAS) transports reducing equivalents between the cytoplasm and the mitochondria and can be important for de novo serine synthesis (Broeks, 2023). Thus, we hypothesized that in serine-deplete H1299 cells, the MAS allows increases in mitochondrial respiration to support cytosolic serine synthesis. To test this, we used CRISPR/Cas9 in H1299 cells to generate isogenic cell lines where either GOT1, MDH1, or GOT2 was deleted (**Supplementary Figure 7A**). Using these knock-out cell lines, we measured proliferation in serine depleted conditions and found that MDH1 and GOT2 knock-out cell lines were more sensitive to serine deprivation compared to non-targeting controls (NTC) (**Supplementary Figure 7B**). We then measured the cell NAD+/NADH ratio in knock-out cells cultured without serine and observed that unlike H1299 NTC and GOT1 KO cells, H1299 MDH1 and GOT2 KO cells did not exhibit elevated NAD+/NADH ratios upon serine depletion (**Supplementary Figure 7C**). Consistently, unlike H1299 NTC cells, H1299 MDH1 and GOT2 KO cells did not demonstrate elevated mitochondrial respiration upon serine depletion (**Supplementary Figure 7D**). This corresponded with H1299 MDH1 and GOT2 KO cells having blunted serine synthesis rates compared to H1299 NTC cells (**Supplementary Figure 7E**). Together, this suggests that MDH1 and GOT2 activity support the process by which mitochondrial NAD+ regeneration is transmitted to the cytoplasm to support serine synthesis.

### The NAD+/NADH ratio influences oxidative citrate synthesis in either serine- or lipid-depleted environments

Oxidative biosynthetic reactions other than serine synthesis can also be constrained by the NAD+/NADH ratio. For example, cancer cells deprived of environmental lipids increase oxidative citrate production, and we have previously found that citrate synthesis, either through glucose oxidation or glutamine oxidation, is limited by NAD+ availability (Li, 2022) (**Figure 5A, Supplementary Figure 8A**). Thus, we sought to uncover whether the increase in the cell NAD+/NADH ratio by mitochondrial respiration in response to serine withdrawal specifically supports greater serine synthesis or also leads to greater oxidative citrate production. To answer this question, we first assessed labeled citrate production using kinetic U-^13^C-glucose tracing in cells that showed increased NAD+/NADH ratios following serine deprivation. We found that the redox responder cells largely showed elevated M+2 citrate production despite no change in environmental lipid availability (**Figure 5B; Supplementary Figure 8B-E**). In contrast, cells with unaltered NAD+/NADH ratios upon serine depletion did not exhibit higher M+2 citrate production **(Figure 5B; Supplementary Figure 8B, F-H)**. Consistently, redox responder H1299 cells, but not redox non-responder A549 cells, displayed evidence of elevated citrate production from glutamine either through oxidative decarboxylation or reductive carboxylation, both of which require oxidation reactions that can be limited by electron acceptor availability (Hosios and Vander Heiden, 2018; Li, 2022) (**Supplementary Figure 8I, J**). To confirm that elevated M+2 citrate synthesis was driven by an increase in the NAD+/NADH ratio, we treated A549 cells depleted of serine with FCCP and examined glucose-derived citrate synthesis. Indeed, FCCP treatment led to a four-fold increase in M+2 citrate from U-^13^C-glucose in A549 cells depleted of serine whereas M+2 citrate synthesis was less affected by FCCP in H1299 cells (**Figure 5C**). The elevated M+2 citrate production in FCCP-treated A549 cells was dependent on mitochondrial complex I, as rotenone prevented the elevated citrate production associated with FCCP treatment (**Supplementary Figure 9A-C**). To confirm that higher M+2 citrate production following FCCP treatment was related to the NAD+/NADH ratio independent of serine deprivation, we treated A549 cells with increasing doses of AKB and FCCP in serine-replete conditions and find that increasing the NAD+/NADH ratio with either AKB or FCCP led to elevated M+2 citrate synthesis (**Figure 5D**). This argues that changes to the NAD+/NADH ratio regardless of the nutrient environment can impact citrate synthesis.

**Figure 5.**
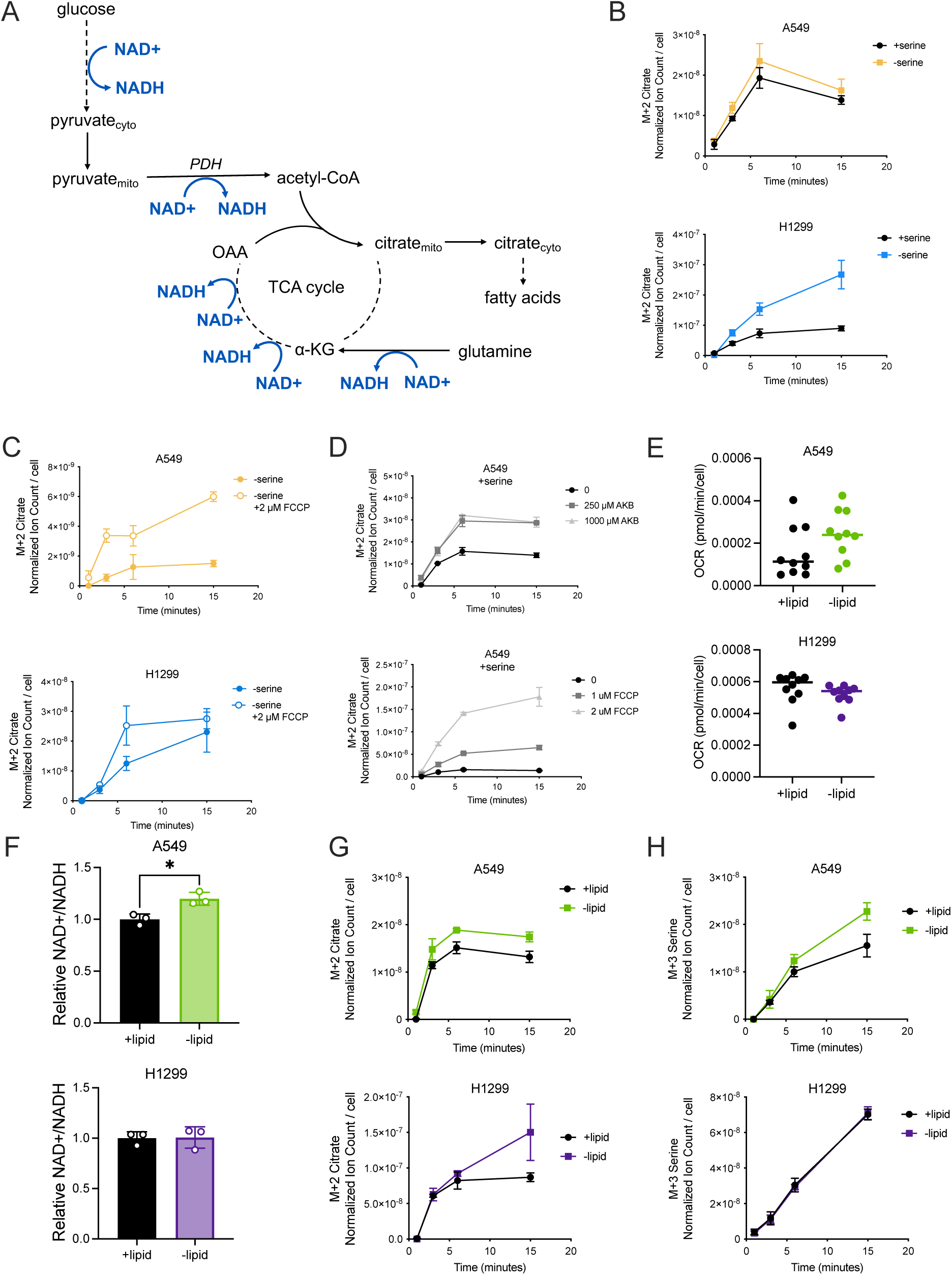
Lipid depletion enhances cell-specific elevation in mitochondrial respiration and the NAD+/NADH ratio, influencing oxidative citrate and serine synthesis. **(A)** Schematic depicting glucose and glutamine oxidative routes for synthesizing citrate, highlighting NAD+ requiring oxidation reactions. **(B)** Citrate labeled (M+2) from U-^13^C-glucose over time in A549 and H1299 cells cultured with or without serine for 24 hours prior to U-^13^C-glucose exposure. Citrate levels are normalized to internal norvaline standard and cell number measured in indicated conditions, n=3. **(C)** Citrate labeled (M+2) from U-^13^C-glucose over time in A549 and H1299 cells cultured without serine and with or without 2 μM FCCP for 24 hours prior to U-^13^C-glucose exposure. Citrate levels are normalized to internal norvaline standard and cell number measured in indicated conditions, n=3. **(D)** Citrate labeled (M+2) from U-^13^C-glucose over time in A549 cells cultured with serine and with indicated treatment conditions for 24 hours prior to U-^13^C-glucose exposure. Citrate levels are normalized to internal norvaline standard and cell number measured in indicated conditions, n=3. **(E)** Oxygen consumption rate (OCR) in A549 and H1299 cells cultured with or without lipids for 24 hours. Values are averages of three repeat measurements, n=10. Green indicates lipid redox responder cells. Purple indicates lipid redox non-responder cells. **(F)** Relative NAD+/NADH ratios of A549 and H1299 cells cultured with or without lipids for 24 hours. NAD+/NADH ratios are normalized to the NAD+/NADH ratio of the corresponding cell type cultured with lipids, n=3. P-values were calculated by unpaired Student’s t-test, *p<0.05. **(G)** Citrate labeled (M+2) from U-^13^C-glucose over time in A549 and H1299 cells cultured with or without lipids for 24 hours prior to U-^13^C-glucose exposure. Citrate levels are normalized to internal norvaline standard and cell number measured in indicated conditions, n=3. **(H)** Serine labeled (M+3) from U-^13^C-glucose over time in A549 and H1299 cells cultured with or without lipids for 24 hours prior to U-^13^C-glucose exposure. Serine levels are normalized to internal norvaline standard and cell number measured in indicated conditions, n=3. Data shown for all panels are means ± SD.

We next asked whether increasing mitochondrial respiration in response to low extracellular serine was specific to serine deprivation or represented a general response to any nutrient deprivation that increased NAD+ demand, such as lipid deprivation (Li, 2022). To examine this, we depleted A549 and H1299 cells of extracellular lipids by culturing cells in media with lipid-depleted serum (Hosios, 2018) and compared oxygen consumption rates to those of cells cultured in media with lipid-containing serum. Surprisingly, we find that A549 cells, which do not increase mitochondrial respiration in response to serine depletion, showed a trend towards a mildly elevated oxygen consumption rate upon lipid depletion (**Figures 5E**). On the other hand, H1299 cells do not increase oxygen consumption in lipid depleted conditions (**Figure 5E**). We next measured the NAD+/NADH ratio of A549 and H1299 cells depleted of environmental lipids. Consistent with the changes in oxygen consumption, A549 cells displayed an increased NAD+/NADH ratio while H1299 cells did not alter the NAD+/NADH ratio upon lipid deprivation (**Figure 5F**). Interestingly, both serine and citrate produced via glucose oxidation was elevated in lipid depleted A549 cells in comparison to H1299 cells, which exhibited unchanged serine synthesis in response to environmental lipid depletion (**Figure 5G-H, Supplementary Figure 10**). These findings suggest that the NAD+/NADH ratio can be a determinant of both oxidative citrate and serine synthesis, regardless of environmental demand for the nutrient. Additionally, the different metabolic responses to lipid and serine deprivation between cancer cell types suggests that different nutrient environments impact mitochondrial respiration through distinct processes to influence the NAD+/NADH ratio.

### Lipid depletion can enhance proliferation in serine deprived conditions

Our finding that both serine and oxidative citrate synthesis can be impacted by levels of a different nutrient than that produced by each pathway raised the possibility that the NAD+/NADH ratio coordinates biomass synthesis in response to the availability of multiple oxidized nutrients. Specifically, we hypothesized that changes to the NAD+/NADH ratio caused by one oxidized nutrient depletion could influence the synthesis of, and in turn the dependency on, another oxidized nutrient for proliferation. To begin testing this, we first examined how mitochondrial respiration and the NAD+/NADH ratio were impacted by dual lipid and serine depletion compared to single nutrient depletions in both A549 and H1299 cells. We find that dual depletion of serine and lipids in A549 cells led to elevated oxygen consumption that is higher than that observed with serine depletion alone **(Figure 6A)**. Similarly, dual depletion of serine and lipids in H1299 cells led to elevated oxygen consumption compared to that observed with lipid depletion alone **(Figure 6A)**. The resulting oxygen consumption rate corresponded with the cell NAD+/NADH ratio in each condition, which was collectively higher upon dual serine and lipid starvation than either serine depletion in A549 cells or lipid depletion in H1299 cells (**Figure 6B**). We then measured the serine and citrate synthesis rates of cells depleted of both extracellular serine and lipids via kinetic U-^13^C-glucose tracing. Consistent with elevation of the NAD+/NADH ratio after lipid deprivation, A549 cells depleted of both environmental serine and lipids trended towards elevated serine synthesis rates when compared to serine deprivation alone, though was not statistically significant **(Figure 6C).** Moreover, lipid depletion appeared to lead to a greater fraction of total serine derived from glucose in serine depleted A549 cells **(Supplementary Figure 11A)**. In contrast, lipid starvation did not alter serine synthesis in serine deprived H1299 cells (**Figure 6D, Supplementary Figure 11A,B**). Interestingly, in both A549 and H1299 cells, dual serine and lipid deprivation led to a trend toward increased citrate production compared to lipid deprivation alone despite only H1299 cells exhibiting a more oxidized NAD+/NADH upon serine depletion (**Figure 6E,F; Supplementary Figure 11C,D)**. This underscores the multifactorial regulation of glucose oxidation to citrate (Holness, 2003; Novelli, 1950; Patel, 2006; Srere, 1974). We next asked how cancer cell proliferation was impacted by dual serine and lipid depletion compared to single nutrient depletions. Strikingly, proliferation of serine depleted A549 cells was improved with depletion of extracellular lipids (**Figure 6G**), consistent with the elevation in mitochondrial respiration, NAD+/NADH ratio, and a trend towards greater serine synthesis in A549 cells upon dual serine and lipid depletion. Unlike serine depletion, lipid depletion does not cause a major proliferation defect in either A549 or H1299 cells, and dual serine and lipid deprivation did not greatly alter the proliferation of H1299 cells compared to serine depletion alone (**Figure 6H**). This is consistent with lipid deprivation causing no major changes in oxygen consumption, NAD+/NADH ratio, and serine synthesis in H1299 cells. These data argue that the cell NAD+/NADH ratio is modulated in response to multiple nutrient availabilities in a cancer cell-specific fashion. The changes in NAD+/NADH ratio in turn may influence the capacity for serine and citrate synthesis, impacting the proliferation rate of cancer cells in nutrient environments with differing demands for oxidative biomass synthesis. Taken together, we propose a model where environmental nutrient availability can impact mitochondrial respiration based on the specific cancer. Because mitochondrial respiration is a major pathway that regenerates NAD^+^, changes to mitochondrial respiration can alter the cell NAD+/NADH ratio, influencing the activity of major NAD^+^-requiring metabolic reactions such as serine synthesis and citrate synthesis that can be important for proliferation. We further propose that changes to the cell NAD+/NADH ratio can impact all oxidative biosynthetic reactions if the enzyme machinery is present, but that specificity for how the cell NAD+/NADH ratio changes is dependent on both cell-intrinsic factors and cell-extrinsic factors (**Figure 7**).

**Figure 6.**
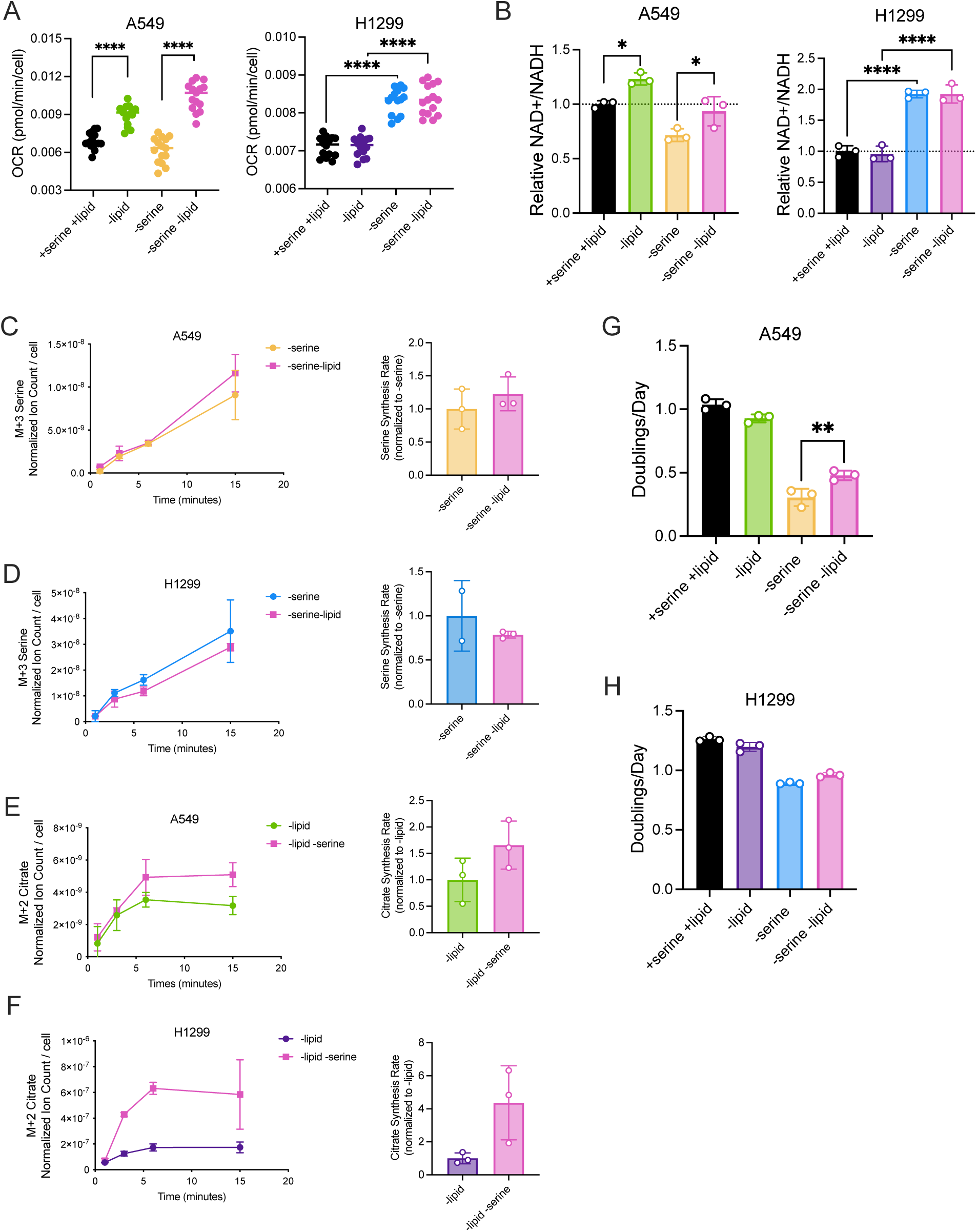
Lipid depletion increases mitochondrial respiration and the NAD+/NADH ratio in a cell-specific manner to influence serine synthesis and proliferation in serine depleted conditions. **(A)** Oxygen consumption rate (OCR) of A549 and H1299 cells cultured with or without serine or lipids for 24 hours. Values are the average of three repeat measurements, n=14-15. P-values were calculated by one-way ANOVA followed by post-hoc Tukey HSD test, ****p<0.001. **(B)** Relative NAD+/NADH ratios of A549 and H1299 cells cultured with or without serine or lipids for 24 hours. NAD+/NADH ratio is normalized to the NAD+/NADH ratio in corresponding cells cultured with serine and lipids, n=3. P-values were calculated by one-way ANOVA followed by post-hoc Tukey HSD test, *p<0.05, ****p<0.001. **(C, D)** Left: Serine labeled (M+3) from U-^13^C-glucose over time in A549 cells (C) and H1299 cells (D) cultured without serine and with or without lipids for 24 hours prior to U-^13^C-glucose exposure. Right: Serine synthesis rates of A549 cells (C) and H1299 cells (D) cultured in indicated conditions. Serine synthesis rates are calculated from labeled serine (M+3) after 1 and 15 minutes of U-^13^C-glucose exposure and normalized to the serine synthesis rates of the corresponding cell line cultured without serine and with lipids. Serine levels are normalized to internal norvaline standard and cell number measured in indicated conditions, n=3. Single data point (H1299, -serine +lipid, t=15 min) removed due to detection error with no signal. (**E,F)** Left: Citrate labeled (M+2) from U-^13^C-glucose over time in A549 cells (E) and H1299 cells (F) cultured without lipids and with or without serine for 24 hours prior to U-^13^C-glucose exposure. Right: Citrate synthesis rates of A549 cells (E) and H1299 cells (F) cultured in indicated conditions. Citrate synthesis rates are calculated from labeled citrate (M+2) after 1 an 15 minutes of U-^13^C-glucose exposure and normalized to the citrate synthesis rates of the corresponding cell line cultured without lipids and with serine. Citrate levels are normalized to internal norvaline standard and cell number measured in indicated conditions, n=3. **(G)** Proliferation rate (doublings per day) of A549 cells cultured with or without serine or lipids for 72 hours, n=3. P-values were calculated by one-way ANOVA followed by post-hoc Tukey HSD test, **p<0.01 **(H)** Proliferation rate (doublings per day) of H1299 cells cultured with or without serine or lipids for 72 hours, n=3. Data shown for all panels are means ± SD.

**Figure 7.**
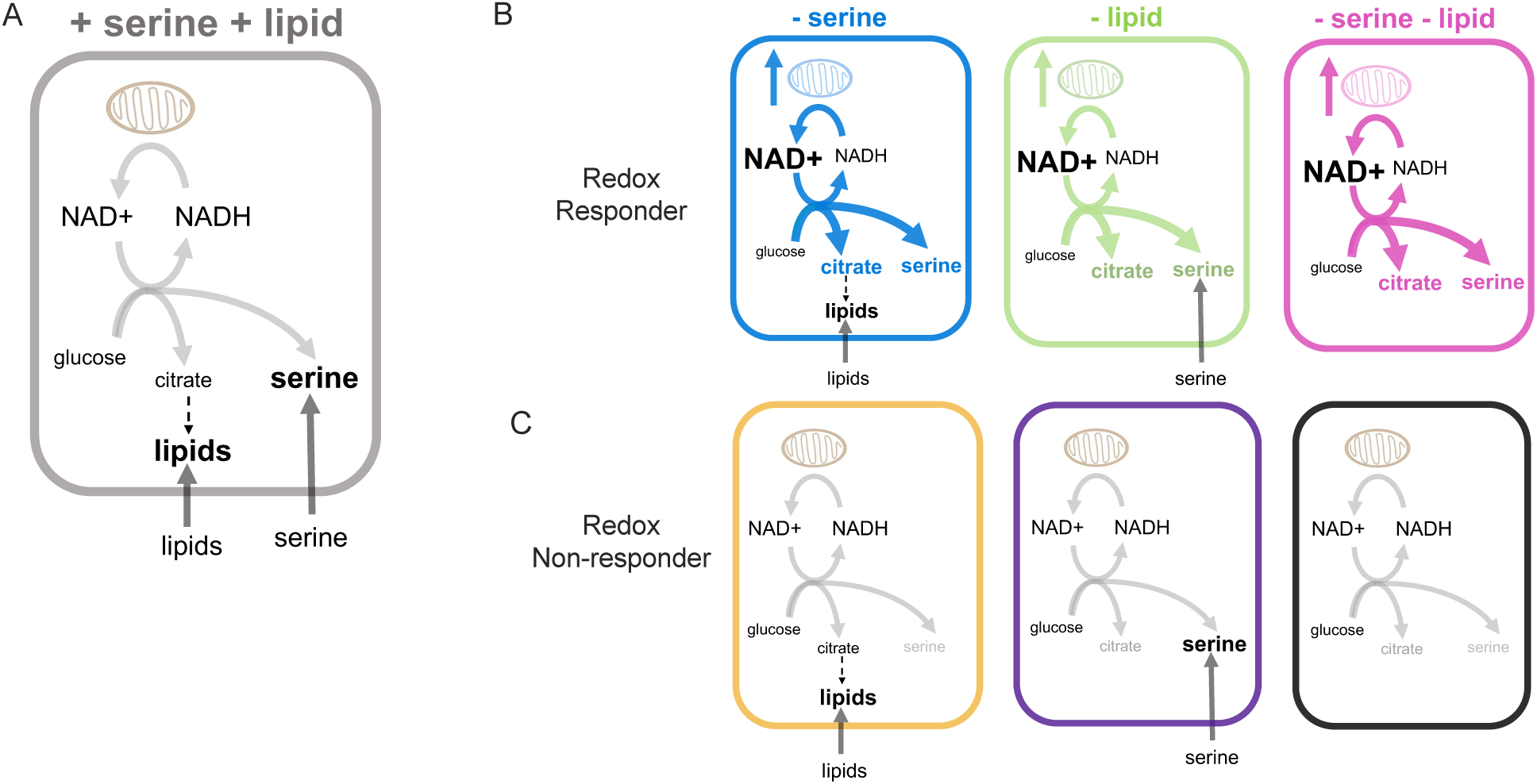
The cell NAD+/NADH ratio is regulated in a cancer cell-specific manner by changes to mitochondrial respiration in response to nutrient availability, impacting oxidative biosynthetic reactions. **(A)** In serine and lipid-replete conditions, cells can acquire environmental serine and lipids to support proliferation. Cells can also oxidize glucose to produce serine, as well as citrate for lipid synthesis. The oxidation reactions involved can be limited by NAD+ availability, which is impacted by mitochondrial respiration. **(B)** In nutrient depleted conditions, the ability to support oxidative reactions is dependent on the mitochondrial response to the nutrient depletion. For redox responders, environmental depletion of serine or lipids increases the cell NAD+/NADH ratio to support both oxidative citrate and serine synthesis. This ability to increase respiration enables proliferation in conditions lacking serine. **(C)** In redox non-responder cells, neither mitochondrial respiration nor the NAD+/NADH ratio increases following nutrient depletion. This limits oxidative synthesis of serine (and citrate), subsequently limiting proliferation in environments that lack serine.

## Discussion

Proliferating cancer cells must acquire sufficient biomass and rely on contributions from both environmental nutrients and de novo synthesis pathways, but what determines which cells can adapt to different nutrient environments that impose constraints on salvage and synthesis is not known. This study highlights that differential cell responses to changes in the NAD+/NADH ratio enable biomass synthesis and determine whether cells can adapt to proliferate in different nutrient conditions. Importantly, this provides one explanation for why single nutrient levels in tissues may not predict the ability of auxotrophy for that nutrient to determine cancer proliferation in a tissue (Abbott, 2026). It also suggests why cancer proliferation in different nutrient environments is controlled by a complex relationship between cell-intrinsic and cell-extrinsic factors. These factors may be coordinated in part through processes that set the NAD+/NADH ratio in cells.

Most prior work to determine what shapes the metabolic dependencies of cell proliferation has focused on factors that govern biosynthetic enzyme expression and activity. For example, genomic and transcriptional analyses of cancer discovered that increased PHGDH expression promotes tumor growth (DeNicola, 2015; Locasale, 2011; Newman, 2017; Possemato, 2011) and provides a proliferative advantage in serine depleted tumor microenvironments (Ngo, 2020; Sullivan, 2019b). Our findings show that enzyme expression alone cannot fully explain nutrient dependencies, as PHGDH expression can be discordant with sensitivity to serine depletion. Rather both the NAD+/NADH ratio and enzyme expression play integral roles in determining cancer proliferation in specific nutrient environments. These data are consistent with biosynthetic gene expression and single nutrient levels being unable to predict the tumor tissue environments where cancer cells can form metastases (Abbott, 2026).

Extensive work has highlighted the importance of cell NAD+/NADH ratio in influencing oxidized biomass production, nutrient utilization, and nutrient dependencies (Baksh, 2020; Bao, 2016; Birsoy, 2015; Broeks, 2023; Christensen, 2014; Diehl, 2019; Garcia-Bermudez, 2018; Gibson, 1976; Krall, 2021; Li, 2022; Sullivan, 2015; Wu, 2024). Yet, the processes that govern the endogenous NAD+/NADH ratio upon shifts in nutrient availability have been largely uncharacterized. We find that regulation of mitochondrial respiration is a major mechanism that dictates the NAD+/NADH ratio in response to environmental nutrient fluctuations. The mitochondrial respiration response to withdrawal of specific nutrients appears to vary across different cancer cells. Some cancer cells increased mitochondrial respiration to serine depletion, while others had minimal responses. We found that in serine redox responder cancer cells, the malate aspartate shuttle (MAS) enzymes MDH1 and GOT2 were required for the elevation in mitochondrial respiration and the NAD+/NADH ratio in response to serine depletion. Thus, differences in the activity of MAS, which can vary greatly across cancer cells (Wang, 2022), may influence the capacity for cancer cells to increase mitochondrial respiration and NAD+ regeneration to support oxidative biosynthetic pathways. Interestingly, serine is an allosteric activator of the glycolytic enzyme pyruvate kinase, which converts phosphoenolpyruvate to pyruvate and generates ATP (Chaneton, 2012). Thus, decreased environment serine availability in addition to differences in pyruvate kinase activity may yield lower glycolytic ATP, resulting in greater mitochondrial respiration in serine redox responder cancer cells. Additionally, it has been recently shown that differences in mtDNA in non-small cell lung cancer cells can lead to defective mitochondrial respiration and imbalanced NAD+/NADH ratios, suppressing serine synthesis and tumor growth (Lopes, 2025). These findings suggest some reasons why respiration may be differentially responsive to serine withdrawal. However, lipid depletion yielded mitochondrial respiration changes in cancer cells that did not respond to serine withdrawal, arguing against respiration defects explaining non-response. No clear common genetic factor or tissue-of-origin was found to align with those cells that do or do not have mitochondrial and redox responses to serine and/or lipid depletion. Further work is needed to understand the mechanisms that enable increased mitochondrial respiration in response to deprivation of different nutrients. One possibility is that differences in ATP consumption following nutrient deprivation contribute to the response. Stimulating ATP consumption can increase mitochondrial respiration and the NAD+/NADH ratio (Bertholet, 2019; Buttgereit, 1995; Fisher-Wellman, 2012; Luengo, 2021; Nobes, 1989; Vander Heiden, 1999). ATP synthase activity can restrain complex I activity to different extents across cancers, modulating the endogenous NAD+/NADH ratio and sensitivity to serine or lipid depletion. While the regulation of complex I, ATP synthesis, and mitochondrial coupling efficiency is complicated and incompletely understood (Alavian, 2014; Gao, 2018; Lapuente-Brun, 2013; Luengo, 2021; Qiu, 2014; Wang, 2022; Yao, 2019; Zhao, 2019), our findings suggest an important connection between the regulation of mitochondrial bioenergetics and specific metabolic vulnerabilities of cancer growth, underscoring an important relationship between environmental nutrient levels and mitochondrial respiration to influence the cell NAD+/NADH ratio.

Efforts to understand the processes that shape cancer cell metabolic dependencies have largely focused on how cancer cells respond to single nutrient changes. However, findings from dual serine and lipid depletion experiments suggest that single nutrient availabilities will be unable to explain cancer metabolic dependencies in some contexts. For example, depleting A549 cells of both extracellular serine and lipids paradoxically improved proliferation relative to serine deprivation alone, consistent with an elevated NAD+/NADH ratio in response to lipid deprivation that can support greater serine synthesis. This suggests that the NAD+/NADH ratio can serve as a metabolic regulatory node that orchestrates biomass synthesis and integrates changes to multiple oxidized biomass components in the environment, providing a framework for understanding why cancer nutrient dependencies vary across tissue microenvironments (Abbott, 2026). Because oxidizing the cell redox state can increase flux through multiple pathways, the impact of cell redox state on biosynthesis may also explain why some cells lose enzyme expression in pathways like serine synthesis, as this could redirect limited resources toward biomass components that can otherwise be scavenged from the environment (Delage, 2010; Pavlova, 2017; Rabinovich, 2015). Better understanding the mechanisms cells use to alter respiration and adjust the NAD+/NADH ratio in response to available nutrients could inform the complex interplay between cell-intrinsic and cell-extrinsic factors that determine cancer metabolic dependencies. This is particularly important to consider when targeting metabolism for cancer treatment. Many newer therapies targeting metabolism have not been successful in part because of metabolic plasticity to nutrient shifts (Amoedo, 2017; Fendt, 2020; Xiao, 2023). Co-targeting mitochondrial function limits metabolic adaptations and may also help predict the tissue nutrient conditions that result in pathway dependencies for specific cancers. Thus, better understanding how the cell NAD+/NADH ratio responds to nutrient levels in different cancers could improve selection of patients for cancer therapies that impact metabolism.

## Material and Methods

### Cell Culture Experiments

Cell lines were maintained in Dulbecco’s Modified Eagle’s Medium (DMEM) without sodium pyruvate (Corning 50-013-PC) supplemented with sodium bicarbonate (Sigma S6014) and 10% heat-inactivated Fetal Bovine Serum (FBS). All cells were cultured at 37°C with 5% CO_2_, tested for mycoplasma contamination regularly, and confirmed negative before experimentation. Cell line information is listed in Supplementary Table 1.

### Proliferation Assays

Cells were plated in six-well plates in 2 ml of DMEM with 10% heat-inactivated FBS at an initial seeding density of 40,000 cells. Cells were permitted to settle overnight and then quantified using a sulforhodamine B (SRB) (Sigma Aldrich 230162) colorimetric assay to measure the starting cell absorbance at the start of the experiment as previously described (Vichai, 2006). For treatment conditions, cells were washed three times with 1 ml of 1X PBS, and 2 ml of treatment media were added to each well. All treatment media was made with 10% dialyzed FBS. Serine-free medium was made by adding a mix of amino acids (Sigma Aldrich) at DMEM concentrations lacking serine and glycine to DMEM with low glucose, without sodium pyruvate, and amino acids (US Biological D9800-13-50L). Glucose and glycine were supplemented for a final concentration for 25 mM and 0.4 mM, respectively. Three days after the initial treatment, cells were quantified using SRB assay.

All SRB measurements are normalized to a blank to correct for background signal. Proliferation rates were calculated with the following formula:

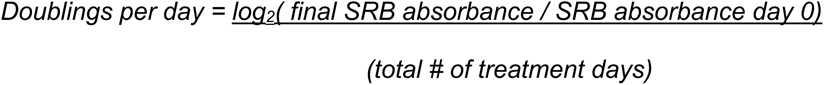

### Cell Number and Viability Measurements

Cells were seeded at an initial density of 2,000 cells per well in a 96-well plate in 200 μl of treatment media. After allowing cells to settle for 1-2 hours, cell number was monitored over time using imaging with the Incucyte Live-Cell Analysis System (Sartorius). Doublings per day were calculated between hour 0 of monitoring and hour 72 of monitoring using the following formula:

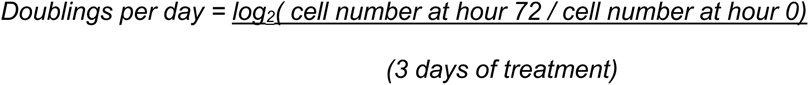

For viability, cells were seeded at an initial density of 2,000 cells per well in a 96-well plate in 200 μl of treatment media with 20 nM Sytox Green (Invitrogen S7020). After 72 hours, the number of live and dead cells present was monitored using imaging with the Incucyte Live-Cell Analysis System (Sartorius).

### NAD+/NADH Measurements

NAD+/NADH measurements were performed using the NAD/NADH-Glo Assay (Promega G9072) with a modified version of manufacturer instructions as previously reported (Sullivan, 2015). Cells were plated at an initial seeding density of 20,000 cells (A549, H1299, HCT116) or 40,000 cells (Calu6, MDA-MB-231). Cells were permitted to settle overnight. The next day, cells were washed once with 1X PBS and then incubated with 2 ml of treatment media for the indicated times prior to preparation of cell extracts. For extraction, cells were washed once in ice cold 1X PBS and extracted in 100 μl ice-cold 1% dodecyltrimethylammonium bromide (DTAB) in 0.2N NaOH diluted 1:1 with 1X PBS. Each sample was flash-frozen in liquid nitrogen and immediately stored at -80°C. To measure NADH, 10 µl of sample was moved to PCR tubes, diluted with 10 µl of DTAB, and incubated at 75 °C for 30 min, where basic conditions selectively degrade NAD+. To measure NAD+, 10 µl of the samples was moved to PCR tubes containing 30 µl DTAB and 20 µl 0.4 N HCl and incubated at 60°C for 15 min, where acidic conditions selectively degrade NADH. Samples were then allowed to equilibrate to room temperature and quenched by neutralizing with 20 µl 0.25 M Tris in 0.2 N HCl (for NADH) or 20 µl 0.5 M Tris base (for NAD+). Manufacturer instructions were followed thereafter to measure NAD+/NADH using a luminometer (Tecan Infinite M200Pro). A standard curve with representative samples was done with each assay to confirm that NAD+ and NADH measurements were in the linear range of detection.

### Glucose Consumption and Lactate Secretion Measurements

Cells were plated at a density of 40,000 cells per well in a six well plate. The next day, cells were washed three times with 1X PBS before 2 ml of treatment media were added to each well. In parallel, 2 ml of treatment media were added to plates without cells. An initial cell number before beginning treatment was quantified via Cellometer (Nexcelom Bioscience). 72 hours later, media was collected from exponentially proliferating cells with a parallel collection of media from wells without cells, which we used to measure initial media concentrations to take into consideration evaporative changes. Cell number was quantified via Cellometer along with the volume of media on cells to consider evaporative changes to metabolite concentrations that may have occurred over 72 hours in culture. Glucose concentrations from media samples were measured on a YSI-2900 Biochemistry Analyzer (Yellow Spring Instruments). Every assay included a glucose (Sigma Aldrich G8270) standard curve (0, 2.5, 5, 10, 20, and 40 mM glucose) and lactate (Sigma Aldrich 71716) standard curve (0, 1.25, 2.5, 5, 10, and 20 mM sodium lactate), made in both control and treatment media. To calculate consumption rate per cell for a given nutrient, fmol of glucose or lactate over time were plotted relative to the area under the curve (fmol / cells · time) of an exponential function fit to the number of cells at the start of the assay and after 72 hours of treatment, as previously described (Hosios, 2016).

### Oxygen Consumption Measurements

An Agilent Seahorse Bioscience Extracellular Flux Analyzer (XFe96) was used to measure oxygen consumption rates (OCR). Cells were plated at 20,000 – 40,000 cells per well in a Seahorse Bioscience 96-well plate in 80 μl of DMEM without pyruvate supplemented with 10% heat-inactivated FBS. Cells were not plated on the perimeter of the plate to avoid edge effect. The following day, cells were washed twice with 180 μl of treatment media supplemented with 10% dialyzed FBS before incubating cells with 180 μl of treatment media for the indicated time prior to OCR data acquisition. For 24 hour treatment periods, treatment media was replaced with 180 μl of fresh treatment media two hours before OCR data acquisition. The day before OCR data acquisition, the XFe96 cartridge was hydrated by submerging the sensor cartridge in 200 μl of sterile water per well of the utility plate. Parafilm was wrapped around the perimeter of the cartridge to avoid evaporation. The XFe96 cartridge was then incubated in a 37°C non-CO_2_ incubator overnight. The next day, at least two hours before data acquisition, the water in the utility plate was replaced with 200 μl of the Agilent XF Calibrant per well and the sensory cartridge was submerged and returned to the 37°C non-CO_2_ incubator until OCR was ready to be measured. After OCR data acquisition, six to eight wells per treatment condition were collected for cell number quantification (Cellometer) to normalize OCR values. Basal OCR was calculated by subtracting residual OCR, the OCR remaining following the addition of rotenone and antimycin A for a final concentration of 2 μM.

### Kinetic U-^13^C-Glucose Isotope Tracing Experiments

Cells were seeded at 150,000 cells per well in a six-well plate in 2ml of DMEM without sodium pyruvate supplemented with 10% heat-inactivated FBS. The following day, cells were washed once with 1ml of 1X PBS and then incubated in 2ml of treatment media supplemented with 10% dialyzed FBS for the indicated treatment time and when metabolic steady state is reached. Two hours before the start of the kinetic isotope tracing, media on cells was exchanged with fresh treatment media with 10mM unlabeled glucose. To measure serine labeling over time in serine depleted conditions, cells and media were extracted together. Before the initiation of the kinetic U-^13^C-glucose isotope tracing, serine-deplete cells were washed three times with 1X PBS and then cultured with 400 μl of tracing medium containing 10 mM of U-^13^C-glucose (Cambridge Isotope Laboratories, CLM-481-0) for the rapid time points. Tracing media was equivalent to treatment media except with 10 mM of U-^13^C-glucose. Following incubation, 1.6ml of extraction buffer containing ice cold 100% HPLC-grade methanol (Sigma Aldrich, 646377) with norvaline (Sigma, N7627) was added onto cells with tracing media for a final sample volume of 2 ml (80% extraction buffer and 20% tracing media) and a final concentration of 1 μg per 400 μl norvaline.

1.6 ml of the lysate was placed into a fresh Eppendorf tube, vortexed at 4°C at maximum speed for ten minutes, and then spun down at 4°C at maximum speed for thirty minutes. The supernatant was collected and dried under nitrogen gas to prepare for metabolic analysis. To measure serine and citrate production in serine replete conditions, only intracellular metabolites were quantified. Before the initiation of kinetic U-^13^C-glucose isotope tracing, serine-replete cells were washed three times with 1X PBS and then cultured with 1 ml of tracing medium containing 10 mM of U-^13^C-glucose (Cambridge Isotope Laboratories, CLM-481-0) for the rapid time points. Tracing media was equivalent to treatment media except with 10 mM of U-^13^C-glucose. Following incubation, cells were rapidly washed twice with ice cold blood bank saline. To extract, 500 μl of extraction buffer containing 80% ice cold HPLC-grade methanol and 20% LC-grade water spiked with 1 μg per 400 μl norvaline were added onto cells. Cells were then lysed as previously described above.

### Gas Chromatography-Mass Spectrometry (GC-MS) Polar Metabolite Measurements

Dried samples were derivatized by adding 16 μl of methoxamine reagent (Thermo Fisher, TS-45950) and incubated for an hour at 37°C followed by the addition of 20 μl of *N*-*tert*-butyldimethylsilyl-*N*-methyltrifluoroacetamide with 1% *tert*-butyldimethylchlorosilane (Sigma 375934). Samples were then incubated for two hours at 60°C. Afterwards, samples were centrifuged at maximum speed for ten minutes, and 20 μl of supernatant was used for analysis. Following derivatization, samples were analyzed using a DB-35MS column (30 m × 0.25 mm i.d. × 0.25 μm, Agilent J&W Scientific) in an Agilent 7890 gas chromatograph coupled to an Agilent 5975C mass spectrometer (GC–MS). All ion counts were normalized to internal standard norvaline and cell number measured via Cellometer.

### Immunoblotting

Cells were washed with ice cold 1X PBS and lysed in cold RIPA buffer containing Halt Protease and Phosphatase Inhibitor Cocktail (Thermo Scientific 78442). Lysates were clarified by rocking samples for thirty minutes at 4°C and then spun at maximum speed for ten minutes for collection of supernatants. Protein concentration was calculating using the BCA Protein Assay (Pierce, 23225) with BSA as a standard. Lysates were resolved by SDS-PAGE using NuPAGE 4-12% Bis-Tris Protein Gels (Thermo Fisher Scientific) and run at 100V. Proteins were transferred onto nitrocellulose membranes via wet transfer at 100V. Membranes were blocked in 5% BSA in 1x TBST before incubating membranes with primary antibodies at 4°C overnight. The next day, membranes were washed three times with 1x TBST at room temperature and then incubated in secondary antibodies for one hour at room temperature. The primary antibodies used were PHGDH (Sigma Aldrich, HPA021241 [1:1000]), PSAT1 (Abcam, ab96136 [1:1000]), PSPH (Abcam, ab96414 [1:1000]), HSP90 (Cell Signaling Technology, 4874 [1:1000]), Vinculin (E1E9V) XP® (Cell Signaling Technology, 13901 [1:1000]), GOT1 (Cell Signaling Technology, 34423S [1:1000], MDH1 (MDHC H-6, Santa Cruz Biotechnology, sc-166879 [1:1000]), GOT2 (Cell Signaling Technology, 71692S [1:1000]), FLAG (Cell Signaling Technology, 14793 [1:1000]). The secondary antibodies used were anti-rabbit IgG horseradish peroxidase-linked antibody (1:4000 dilution; Cell Signaling Technologies, 7074S) or anti-mouse IgG horseradish peroxidase-linked antibody (1:4000, 7076S). Protein expression was quantified by using ImageJ analysis.

### Genetic Manipulation of Enzyme Expression in Cells

Cell lines overexpressing either EGFP or human PHGDH were generated via lentiviral infection. pLJM1-EGFP was obtained from Addgene (Addgene 19319). pLJM1-PHGDH was constructed using pLJM1-Empty (Addgene 91980) as a backbone. pLJM1-Empty was digested using NheI and EcoRI and gel purified. Human PHGDH cDNA was amplified from pLHCX-PHGDH (PMID: 31598584) with the following oligonucleotide primers, where the capitalized sequence denotes

PHGDH homology: PHGDH F: agtgaaccgtcagatccggctagcgccaccATGGCTTTTGCAAATCTGCG

PHGDH R: tactgccatttgtctcgaggtcgagaattcTTAGAAGTGGAACTGGAAGGCT

PHGDH insert was gradient PCR amplified using Phusion High-Fidelity DNA polymerase (NEB M0530) from 60 to 70°C and gel purified. Digested pLJM1-Empty and amplified PHGDH insert were assembled via Gibson Assembly, and the resulting plasmid was transformed, and validated by Sanger sequencing. Lentiviral production was done by transfecting constructs into LentiX293T cells with Mirus Transit293T (Mirus Bio MIR2700) following manufacturer’s protocol. Briefly, 800,000 cells were plated per well in a 6 well plate in DMEM media with 10% heat-inactivated FBS. The next day, cells were transfected with 1.6 μg vector (either pLJM1-EGFP or pLJM1-PHGDH), 800 ng of pMDLg (Addgene 12251) packaging plasmid, 400 ng of pMD2.G (Addgene 12259) envelope plasmid, and 400 ng of pRSV-REV (Addgene 12253) in OptiMEM (Thermo Fisher Scientific 31985070). DNA transfection mix was incubated with cells for 48 hours before harvesting virus and filtering through a 0.45 μm polyethersulfone membrane. Sub-confluent cells were infected with 1 ml of virus-containing media in each well of a 6 well plate with polybrene (8 μg/ml, EMD Millipore TR-1003-G). 24 hours after infection, selection with 1 μg/mL puromycin (Sigma Aldrich P7255) began. All experiments with generated cell lines expressing PHGDH were conducted on polyclonal cell populations.

cyto*Lb*NOX-FLAG cDNA and mito*Lb*NOX-FLAG cDNA were PCR amplified from pUC57-*Lb*NOX or pUC57-mito*Lb*NOX, respectively. pUC57-*Lb*NOX (Addgene 75285) and pUC57-mito*Lb*NOX (Addgene 74448) were a gift from Vamsi Mootha. The PCR products were subsequently cloned into pLJM1 using Gibson Assembly (NEB E2611S). pLJM1-EGFP was a gift from David Sabatini (Addgene 19319). Cells were then transduced with lentivirus containing pLJM1-*Lb*NOX-FLAG, pLJM1-mito*Lb*NOX-FLAG, or pLJM1-EGFP. The infected cells were selected in 1 μg/mL puromycin (Sigma Aldrich P7255). All experiments with generated cell lines involving *Lb*NOX were conducted on a polyclonal cell population. GOT1, GOT2, and MDH1 Knockout H1299 cells were generated by using lentiCRISPRv2 (Addgene 52961). LentiCRISPRv2 was a gift from Feng Zhang. Guide RNAs (gRNAs) for each gene of interest were designed by CRISPick (https://portals.broadinstitute.org/gppx/crispick/public) and cloned into lentiCRISPRv2. The guide sequences used are as follows:

NTC: GTGTAGTTCGACCATTCGTG

GOT1: GGAGGTGTGCAATCTTTGGG

GOT2: CGGACGCGGGTCCACTCCCG

MDH1: CGTCCAGGACACCCATCATG

Cell lines were generated by transducing cells with viral particles containing lentiCRISPRv2 followed by selection with 1 µg/mL puromycin (Sigma Aldrich P7255). Single cells were plated into individual wells of 96 well plates in RPMI medium to obtain single clone knock out cells. Following expansion, immunoblotting was used to confirmed complete knockout.

### Statistics and Reproducibility

All statistical tests using experimental data were performed with Prism 10 software. No statistical method was used to predetermine sample sizes. Samples sizes were chosen based on pilot experiments using three or more technical replicates. The experiments were not randomized and the investigators were not blinded during experiments nor data analysis.

## Acknowledgements

We thank all members of the Vander Heiden lab for helpful discussions, Emma Dawson, Grace Wolczanski, and Alejandra Rosario-Crespo for technical assistance and Jessica Spinelli for assistance with using Seahorse equipment during the COVID-19 pandemic. The authors acknowledge support from F30CA268633 (SMC), MSTP T32GM007753 (SMC), T32GM007287 (SMC, MBM, ASZ, KLA), the Kosciuszko Foundation (SET), T32GM144273 (HC), NSF DGE-1122374 (KLA), and F31CA271787 (KLA), as well as support to MGVH from the Ludwig Center at MIT, the MIT Center for Precision Cancer Medicine, and the NCI (R35CA242379, R01CA259253, P30CA1405141).

## Author Contributions

Conceptualization: SMC, MGVH; Intellectual Discussion: SMC, MBM, MGVH; Proliferation rates: SMC, SET, ASZ, HC; Incucyte: SMC, MBM; Viability: SMC, MBM; Kinetic Isotope Tracing: SMC, SET, MBM, HC; Immunoblotting: SMC, SET; NAD+/NADH measurements: SMC, SET; Oxygen consumption: SMC; PHGDH overexpression constructs: KLA; PHGDH overexpression cell line generation: SMC; GOT1, GOT2, and MDH1 KO generation: RC; Writing: SMC and MGVH; Editing: SMC, MBM, SET, ASZ, KLA, MGVH; Funding Acquisition: MGVH

## Supplementary Material

**Supplementary Figure 1.**
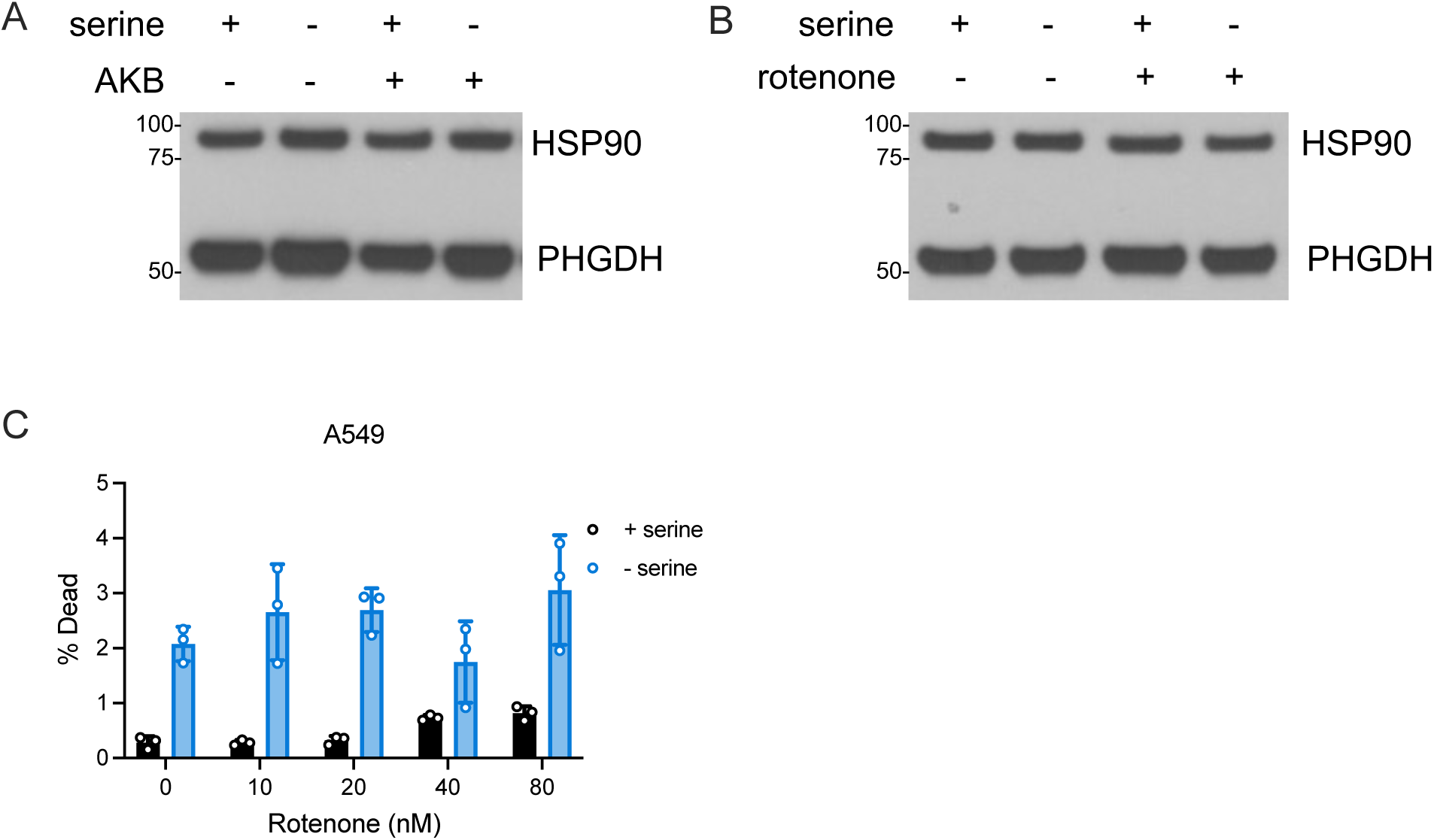
Altering NAD+/NADH with α-ketobutyrate (AKB) or rotenone does not change PHGDH protein levels and minimally changes cell viability. **(A,B)** Immunoblots assessing PHGDH protein levels in A549 cells cultured with or without serine, 1 mM AKB (A), or 20 nM rotenone (B), as indicated for 24 hours. HSP90 expression is shown as a loading control. **(C)** Percent of A549 cells dead, measured using dye exclusion 72 hours after exposure to the indicated concentrations of rotenone in serine-replete or serine depleted conditions, n=3. Serine depletion led to a statistically significant increase in cell death compared to serine-replete conditions at all concentrations of rotenone with the exception of 40 nM rotenone, p<0.05. There was no statistical difference in cell death between 0 and 20 nM rotenone in either serine-replete or serine depleted conditions. P-values were calculated using unpaired Student’s t-test. Data shown are means ± SD.

**Supplementary Figure 2.**
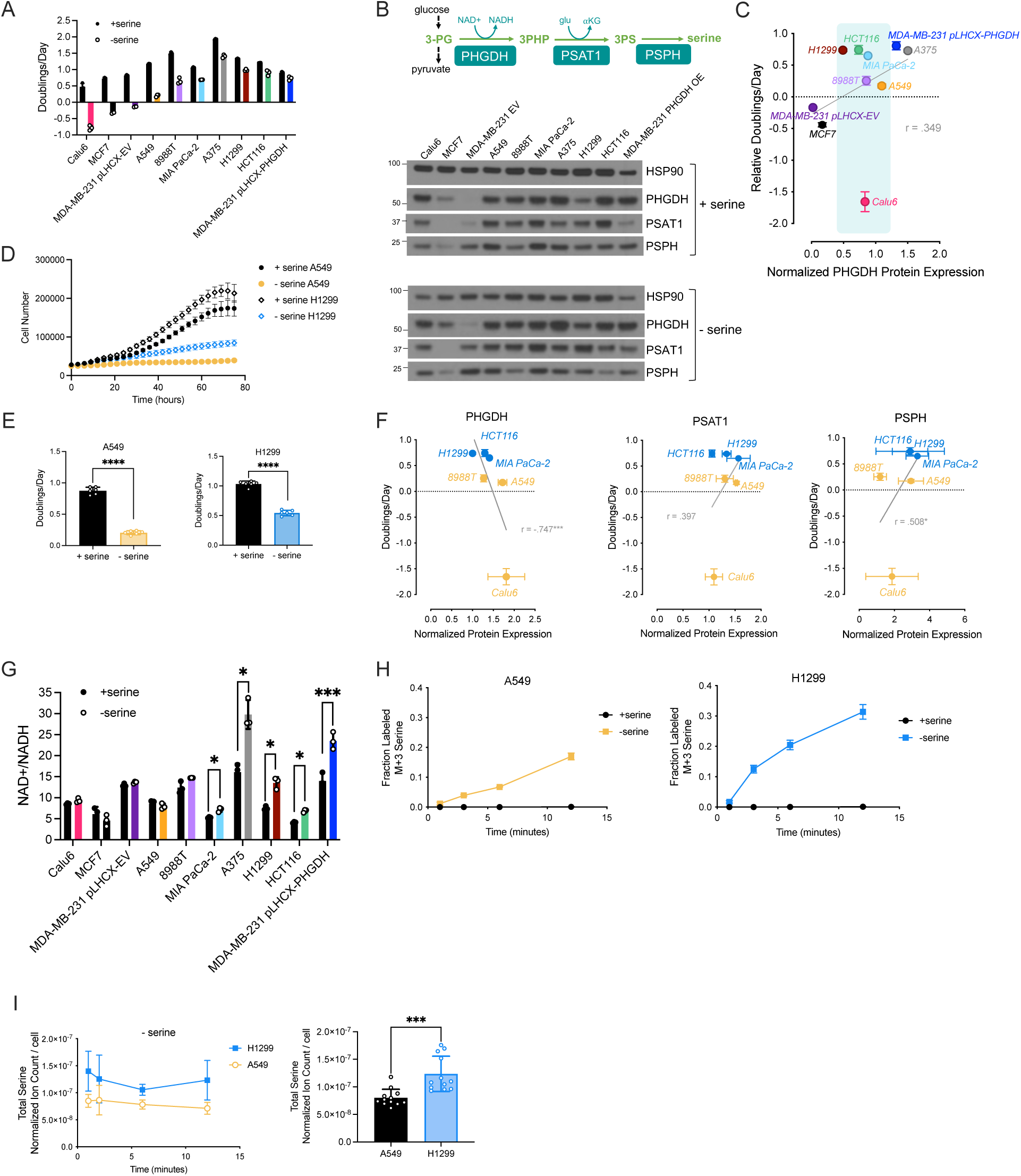
Relationship between PHGDH protein expression, cell NAD+/NADH ratio, and cell proliferation rate following serine deprivation. **(A)** Proliferation rates (doublings per day) of cells cultured with or without serine for 72 hours, n=3. **(B)** Above: Schematic of the serine synthesis pathway. Below: Immunoblot assessing protein levels of PHGDH, PSAT1, and PSPH in the indicated cells cultured with or without serine for 24 hours. HSP90 protein expression is shown as a loading control. These data are representative of three independent experiments. Abbreviations – 3-PG: 3-phosphoglycerate, PHGDH: phosphoglycerate dehydrogenase, 3PHP: 3-phosphohydroxypyruvate, glu: glutamate, αKG: α- ketoglutarate, PSAT1: phosphoserine aminotransferase 1, 3PS: 3-phosphoserine, PSPH: phosphoserine phosphatase**. (C)** Quantification of PHGDH protein expression for the indicated cells under serine depleted conditions (normalized to HSP90 protein expression) from three biological replicates, correlated with the corresponding relative proliferation rate of indicated cells under serine depleted conditions based on data presented in (A). Cell lines enclosed in the light blue shaded box indicate cancer cell lines with serine synthesis enzyme expression that could not fully predict proliferation in the absence of serine that we chose to examine further. Pearson correlation coefficient calculated by simple linear regression. **(D)** Cell number over time for either A549 cells (yellow) or H1299 cells (blue) in either serine-replete or serine depleted conditions, n=6. **(E)** Calculated doublings per day from the data in (D) for the indicated cells and conditions, n=6. P-values were calculated using unpaired Student’s t-test, ****p<0.001. **(F)** Correlation between the protein expression of serine synthesis enzymes PHGDH, PSAT1, and PSPH across three biological replicates and proliferation rate in serine depleted conditions for the indicated cell lines, n=3. Pearson correlation coefficient and P-values were calculated by simple linear regression, *p<0.05, ***p<0.005. **(G)** Unnormalized NAD+/NADH ratios measured for the indicated cell lines cultured with or without serine for 24 hours, n=3 per each cell line. P-values were calculated by unpaired Student’s t-test, *p<0.05, ***p<0.005. **(H)** Fraction of total serine labeled from U-^13^C-glucose over time in A549 and H1299 cells that were cultured with or without serine for 24 hours prior to U-^13^C-glucose exposure as indicated, n=3. **(I)** Left: Total intracellular serine levels over time from the kinetic tracing with A549 and H1299 cells cultured with or without serine for 24 hours prior to U-^13^C-glucose exposure as described in (H). Right: Average of total intracellular serine levels from all time points of the kinetic U-^13^C-glucose isotope tracing experiment shown in (H). Serine levels were normalized to internal norvaline standard and cell number. P-values were calculated by unpaired Student’s t-test, ***p<0.005. Data shown for all panels are means ± SD.

**Supplementary Figure 3.**
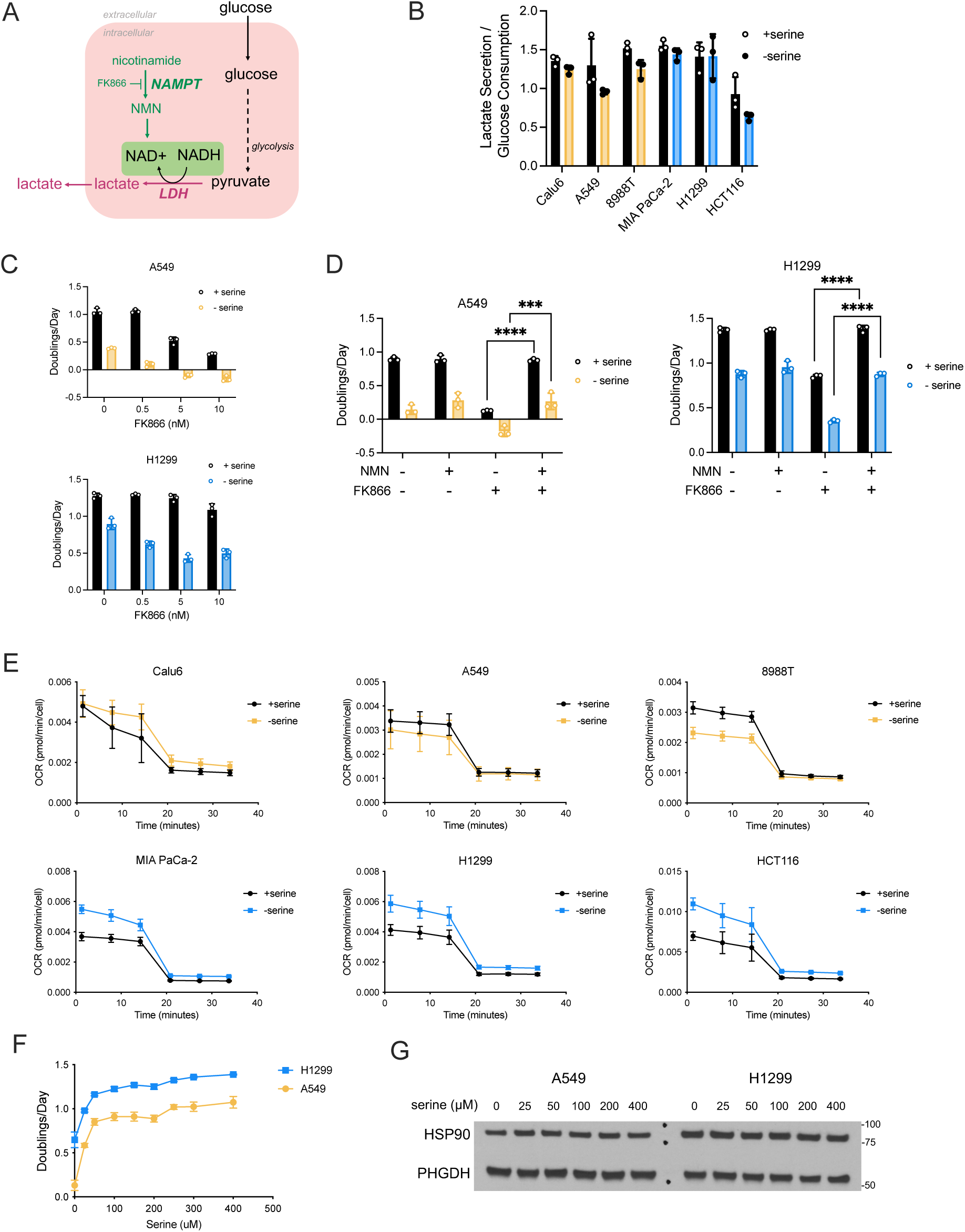
Lactate dehydrogenase activity, reliance on the NAD+ salvage pathway, mitochondrial respiration, and PHGDH protein levels following serine deprivation. **(A)** Schematic depicting the contributions of lactate dehydrogenase (LDH) and the NAD+ salvage pathway to NAD+ generation. Glucose is consumed by cells and converted to pyruvate via glycolysis. LDH can convert pyruvate to lactate by oxidizing NADH to NAD+. In the NAD+ salvage pathway, nicotinamide can be converted to nicotinamide mononucleotide (NMN) by the enzyme nicotinamide phosphoribosyltransferase (NAMPT), which is inhibited by the small molecule FK866. NMN is subsequently converted to NAD+ via NMN adenylyltransferase. **(B)** Lactate secretion rate normalized to glucose consumption rate as measured in the indicated cells cultured with or without serine for 72 hours, n=3. **(C)** Proliferation rate (doublings per day) of A549 (above) and H1299 cells (below) cultured with or without serine and with or without the indicated concentrations of FK866 for 72 hours, n=3. There was a statistically significant difference in the sensitivity of A549 and H1299 cells to FK866 in the serine-replete condition (p<0.001), but not in the serine depleted conditions. P-values were calculated by ANCOVA analysis followed by post-hoc Tukey HSD test. **(D)** Proliferation rate (doublings per day) of A549 (left) and H1299 cells (right) cultured with or without serine and either NMN (100 μM) or FK866 (10 μM) as indicated for 72 hours, n=3. P-values were calculated using unpaired Student’s t-test, ***p<0.005, ****p<0.001. **(E)** Oxygen consumption rate (OCR) of cells cultured with or without serine for 24 hours, n=10-20. Rotenone and antimycin A were injected between the third and fourth OCR measurements to define non-mitochondrial OCR. Data shown are normalized to cell number. **(F)** Proliferation rate (doublings per day) of A549 and H1299 cells cultured in media with the indicated concentrations of serine for 72 hours. Media was replenished every day to avoid serine depletion in lower serine-containing conditions, n=3. **(G)** Immunoblot assessing PHGDH protein levels in A549 or H1299 cells cultured in media with the indicated concentrations of serine for 24 hours. HSP90 expression is shown as a loading control. Data shown for all panels are means ± SD.

**Supplementary Figure 4.**
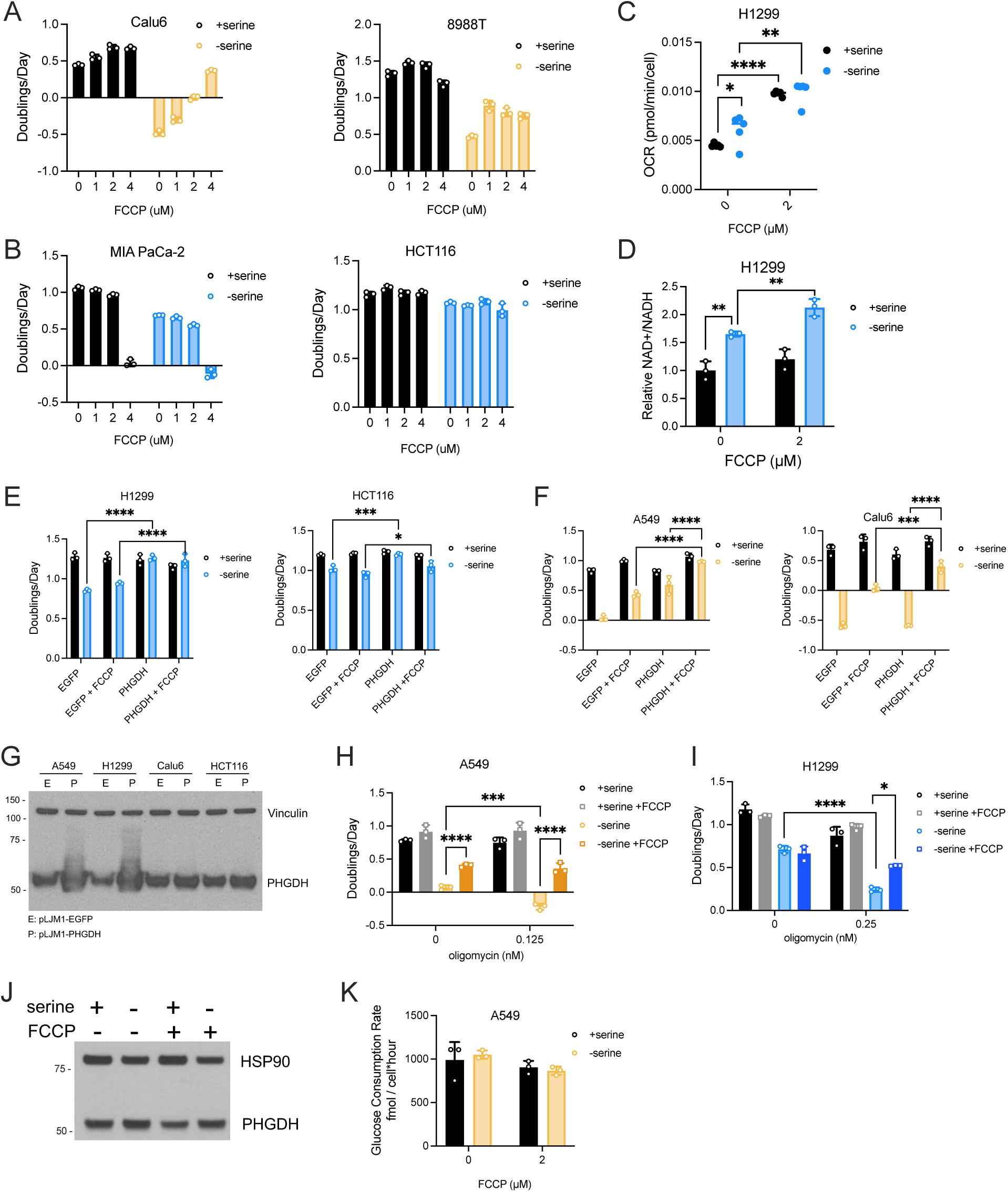
FCCP increases both serine synthesis and proliferation of redox non-responder cells in serine depleted conditions. **(A)** Proliferation rate (doublings per day) of redox non-responder cells Calu6 and 8988T cultured with or without serine and the indicated concentration of FCCP for 72 hours, n=3. Increasing doses of FCCP statistically increased the proliferation of Calu6 cells cultured without serine, p<0.001. P-values were calculated by simple linear regression. All doses of FCCP statistically increased the proliferation of 8988T cells cultured without serine but not in a dose-dependent manner, p<0.001. P-values were calculated by unpaired Student’s t-test. **(B)** Proliferation rate (doublings per day) of redox responder cells MIA PaCa-2 and HCT116 cultured with or without serine and the indicated concentration of FCCP for 72 hours, n=3. **(C)** Mitochondrial oxygen consumption rate (OCR) of H1299 cells cultured with or without serine and the indicated concentration of FCCP for 24 hours, n=5. Values are averages of three repeat measurements. P-values were calculated by unpaired Student’s t-test, *p<0.05, **p<0.01, ****p<0.001. **(D)** NAD+/NADH ratio measured in H1299 cells cultured with or without serine and the indicated concentration of FCCP for 24 hours, n=3. P-values were calculated by unpaired Student’s t-test, **p<0.01. **(E)** Proliferation rate (doublings per day) of redox responder cells H1299 and HCT116 with or without PHGDH overexpression and cultured with or without serine and 2 μM FCCP, as indicated, for 72 hours, n=3. P-values were calculated by two-way ANOVA followed by post-hoc Tukey HSD test, *p<0.05, ***p<0.005, ****p<0.001. **(F)** Proliferation rate (doublings per day) of redox non-responder cells A549 and Calu6 with or without PHGDH overexpression and cultured with or without serine and 2 μM FCCP for 72 hours, n=3. P-values were calculated by two-way ANOVA followed by post-hoc Tukey HSD test, ***p<0.005, ****p<0.001. **(G)** Immunoblot assessing PHGDH protein levels in the indicated cell lines without (pLJM1-EGFP) or with (pLJM1-PHGDH) exogenous PHGDH expression. Vinculin expression is shown as a loading control. **(H,I)** Proliferation rate (doublings per day) of A549 cells (H) and H1299 cells (I) cultured with or without serine, with or without 2 μM FCCP, or the indicated concentration of oligomycin for 72 hours, n=3. P-values were calculated by unpaired Student’s t-test, *p<0.05, ***p<0.005, ****p<0.001. **(J)** Immunoblot assessing PHGDH protein levels in A549 cells cultured with or without serine and with or without 2 μM FCCP for 24 hours. HSP90 expression is shown as a loading control. **(K)** Glucose consumption rates of A549 cells cultured with or without serine and 2 μM FCCP for 72 hours, n=3. Data shown for all panels are means ± SD.

**Supplementary Figure 5.**
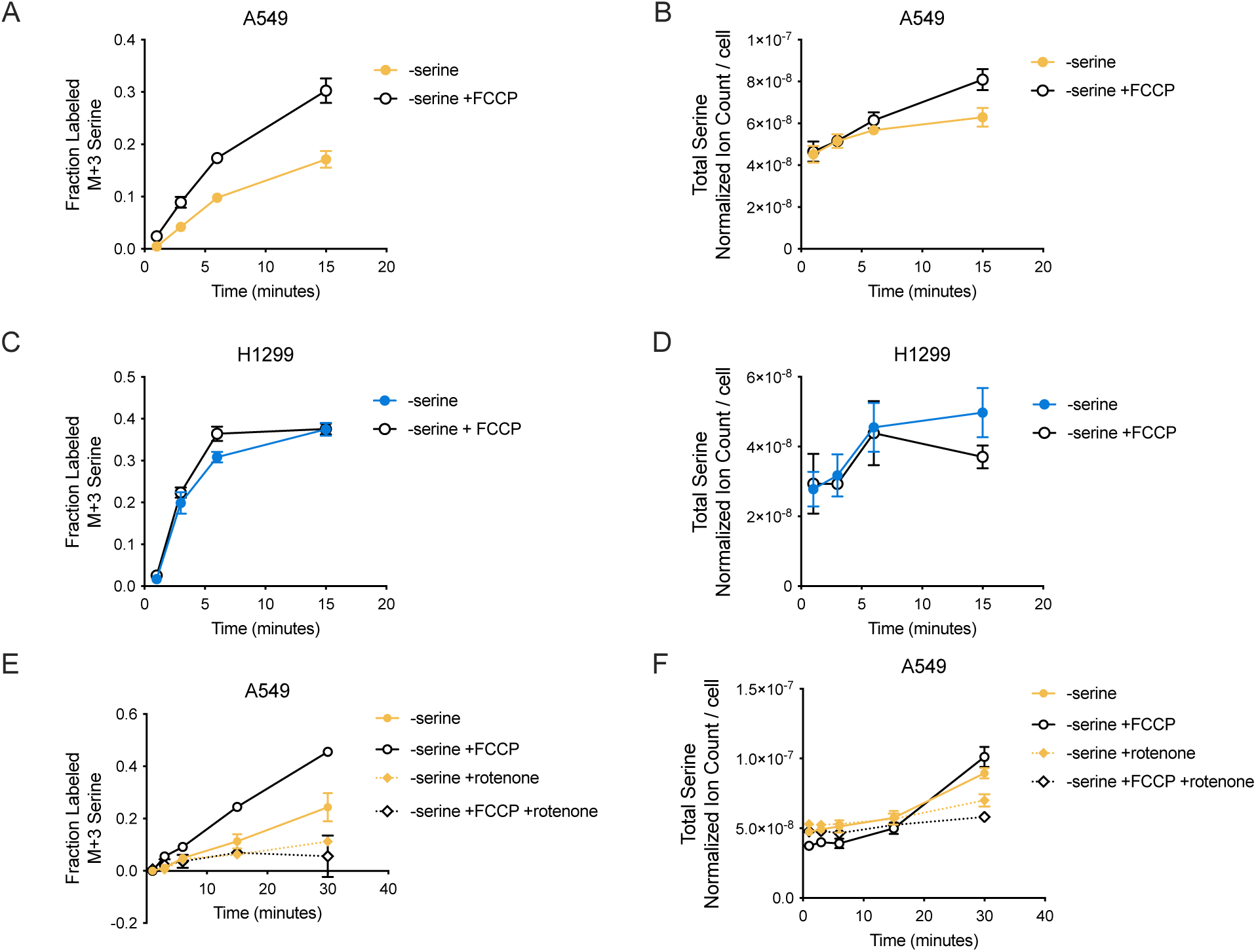
Kinetic tracing of labeled carbon from U-^13^C-glucose into serine following serine withdrawal and FCCP treatment. **(A)** Fraction of total serine labeled (M+3 serine) from U-^13^C-glucose over time in A549 cells cultured without serine, and with or without 2 μΜ FCCP, for 24 hours prior to U-^13^C-glucose exposure as indicated, n=3. **(B)** Total serine measured in A549 cells cultured without serine and with or without 2 μΜ FCCP for each time point shown in (A), n=3. **(C)** Fraction of total serine labeled (M+3 serine) from U-^13^C-glucose over time in H1299 cells cultured without serine and with or without 2 μΜ FCCP for 24 hours prior to U-^13^C-glucose exposure as indicated, n=3. **(D)** Total serine measured in H1299 cells cultured without serine and with or without 2 μΜ FCCP for each time point shown in (C), n=3. **(E)** Fraction of total serine labeled (M+3 serine) from U-^13^C-glucose over time in A549 cells cultured without serine and with or without 2 μΜ FCCP and 20 nM rotenone for 24 hours prior to U-^13^C-glucose exposure as indicated, n=3. **(F)** Total serine measured in A549 cells cultured without serine and with or without 2 μΜ FCCP and 20 nM rotenone for each time point shown in (E), n=3. All serine measurements were normalized to internal norvaline standard and cell number. Data shown for all panels are means ± SD.

**Supplementary Figure 6.**
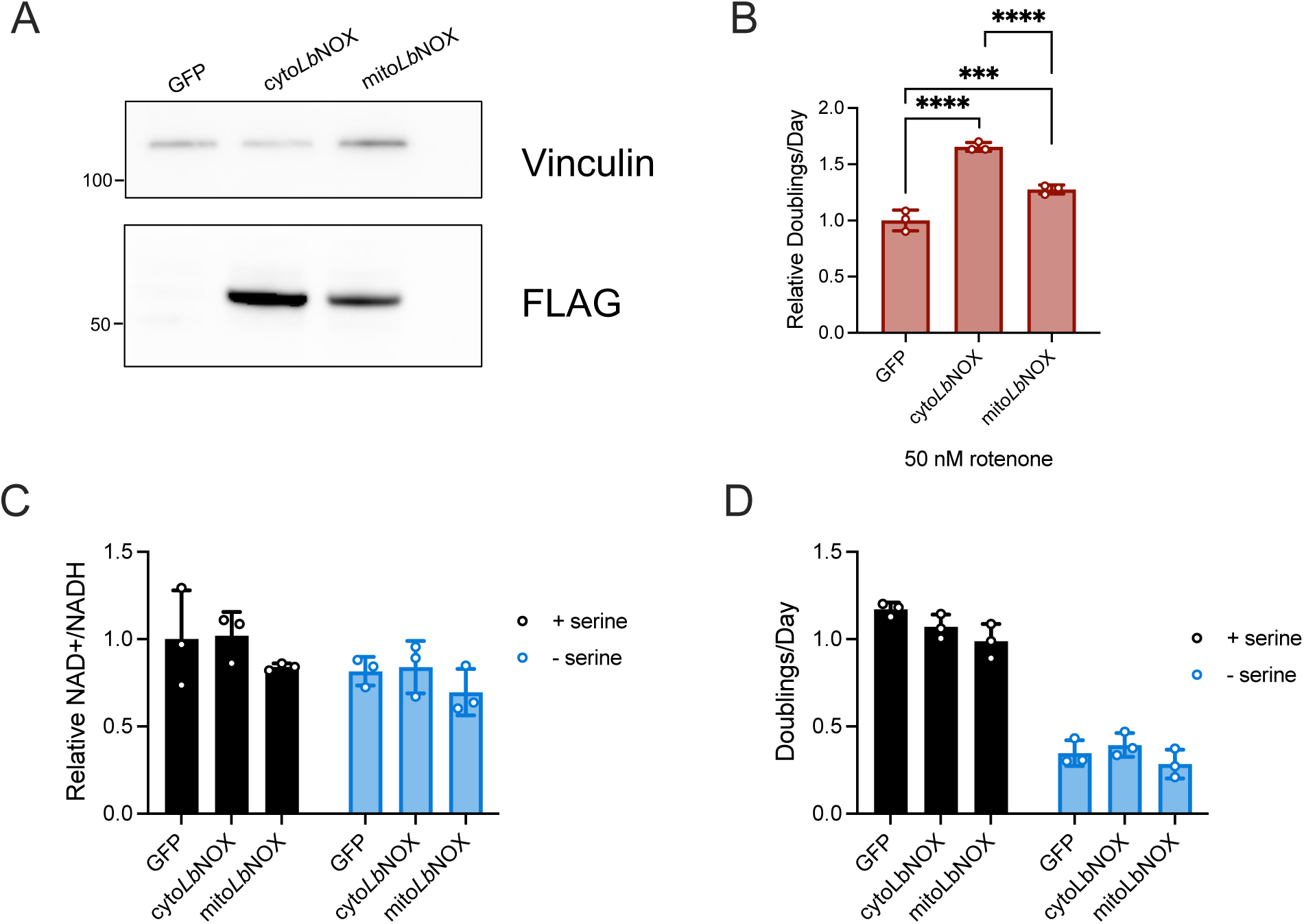
Cytosolic or mitochondrial *Lb*NOX expression does not change the cell NAD+/NADH ratio or proliferation of A549 cells cultured without serine. **(A)** Immunoblot for expression of FLAG-tagged cyto*Lb*NOX or mito*Lb*NOX as indicated. GFP-expressing cells were used as a control. Vinculin expression is shown as a loading control. **(B)** Relative proliferation rate (doublings per day) of A549 cells expressing either cyto*Lb*NOX or mito*Lb*NOX and treated with 50 nM rotenone for 72 hours, n=3. P-values were calculated using one-way ANOVA followed by post-hoc Tukey HSD test, ***p<0.005, ****p<0.001. **(C)** Relative NAD+/NADH ratios of cells expressing either GFP, cyto*Lb*NOX, or mito*Lb*NOX and cultured with or without serine for 24 hours. NAD+/NADH ratios were normalized to the GFP control cells cultured with serine, n=3. **(D)** Proliferation rate (doublings per day) of cells expressing either GFP, cyto*Lb*NOX, or mito*Lb*NOX and cultured with or without serine for 72 hours, n=3. Data shown for all panels are means ± SD.

**Supplementary Figure 7.**
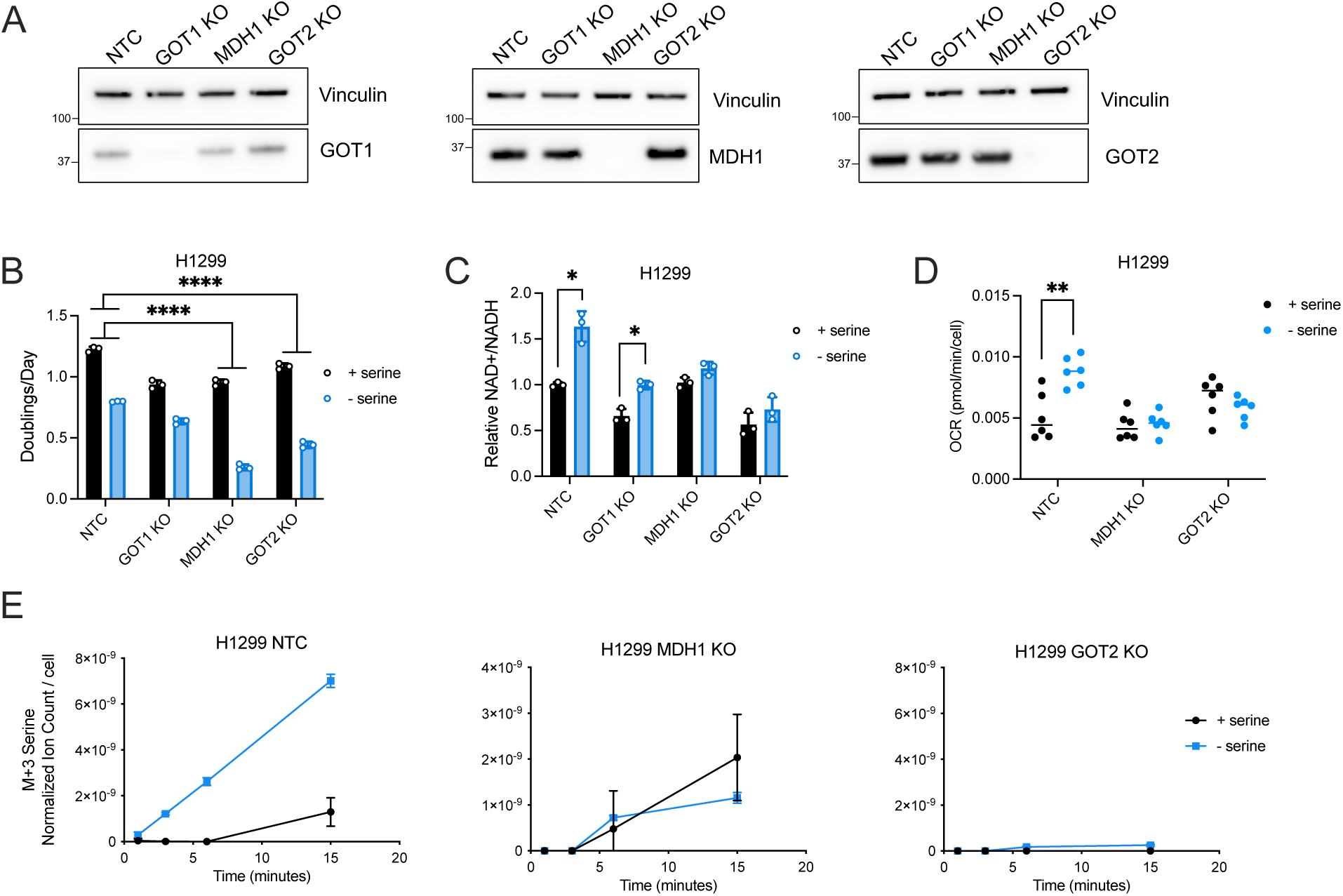
MDH1 and GOT2 support the increase in serine synthesis observed in H1299 cells following serine withdrawal. **(A)** Immunoblots for GOT1 (left), MDH1 (middle), and GOT2 (right) in control cells (NTC, non-targeting control) or cells with knockout (KO) of GOT1, MDH1, or GOT2 as indicated. Vinculin expression is shown as a loading control. **(B)** Proliferation rate (doublings per day) of the indicated cells from (A) cultured with or without serine for 72 hours, n=3. P-values were calculated by nested ANOVA to compare sensitivity to serine depletion between cell lines, ****p<0.001. **(C)** Relative NAD+/NADH ratios of the indicated cells cultured with or without serine for 24 hours, n=3. NAD+/NADH ratios were normalized to H1299 NTC cells cultured with serine. P-values were calculated using unpaired Student’s t-test, *p<0.05. **(D)** Oxygen consumption rate (OCR) of the indicated cells cultured with or without serine for 24 hours, n=6. P-values were calculated by unpaired Student’s t-test, **p<0.01. **(E)** Serine labeled from U-^13^C-glucose (M+3) measured over time from the indicated cells cultured with or without serine for 24 hours prior to U-^13^C-glucose exposure, n=3. Serine levels were normalized to internal standard norvaline and cell number. Data shown for all panels are means ± SD.

**Supplementary Figure 8.**
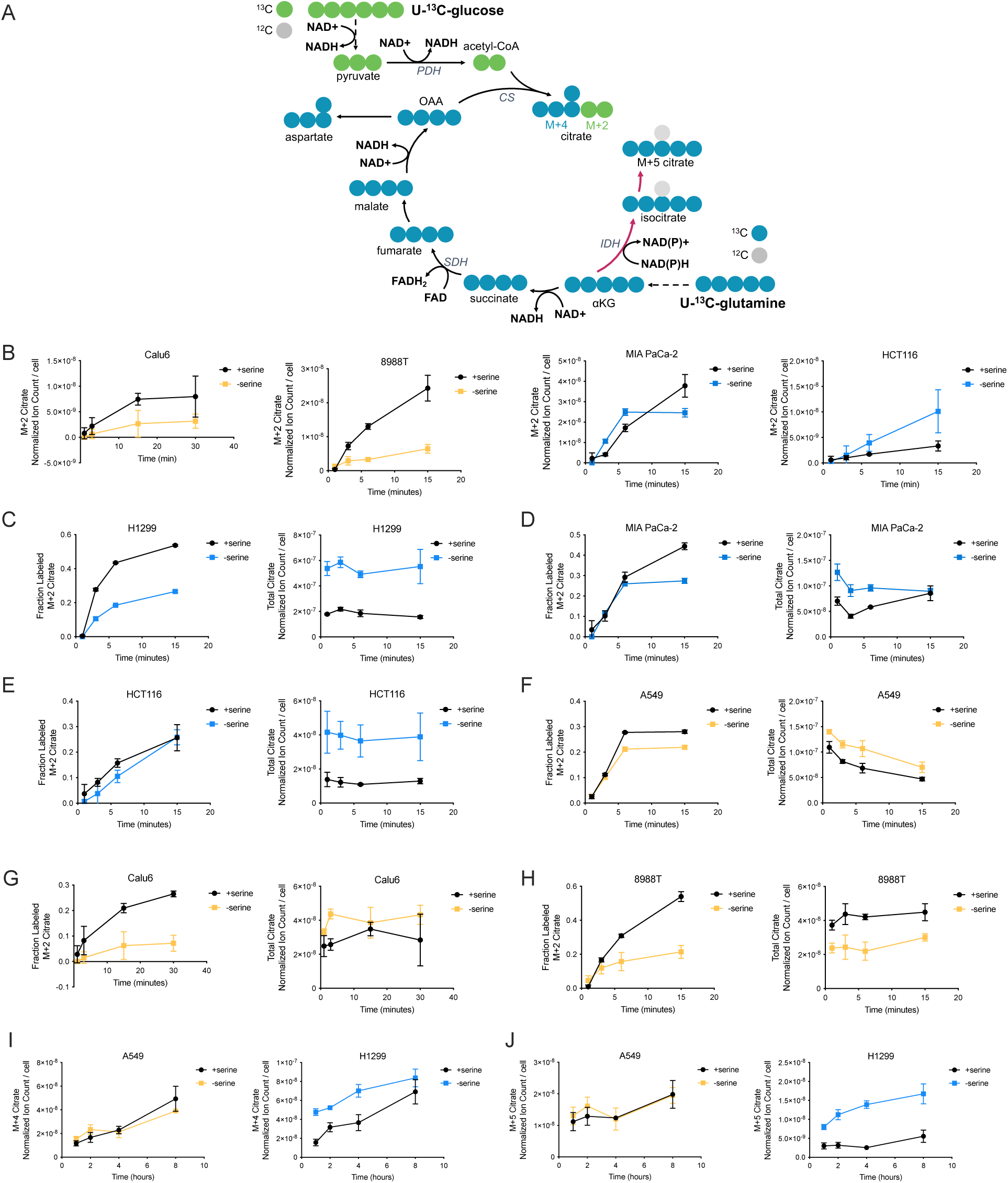
Kinetic tracing of labeled carbon from U-^13^C-glucose and U-^13^C-glutamine into citrate in cells cultured with or without serine. **(A)** Schematic depicting isotopically-labeled carbon contribution to citrate from U-^13^C-glucose and U-^13^C-glutamine. Abbreviations: PDH – pyruvate dehydrogenase, CS – citrate synthase, IDH – isocitrate dehydrogenase, SDH – succinate dehydrogenase. **(B)** Citrate labeled (M+2) from U-^13^C-glucose over time in Calu6, 8988T, MIA PaCa-2, and HCT116 cells cultured with or without serine for 24 hours prior to U-^13^C-glucose exposure, n=3. **(C-H)** Left of each figure panel: Fraction of citrate labeled (M+2) from U-^13^C-glucose over time in indicated cells cultured with or without serine for 24 hours, n=3. Right of each figure panel: Total citrate levels in the indicated cells cultured with or without serine for 24 hours for each time point shown in left panel, n=3. **(I)** Citrate labeled (M+4) from U-^13^C-glutamine over time in A549 and H1299 cells cultured with or without serine for 24 hours, n=3. **(J)** Citrate labeled (M+5) from U-^13^C-glutamine over time in A549 and H1299 cells cultured with or without serine for 24 hours, n=3. All citrate measurements were normalized to internal norvaline standard and cell number measured in indicated conditions. Data shown for all panels are means ± SD.

**Supplementary Figure 9.**
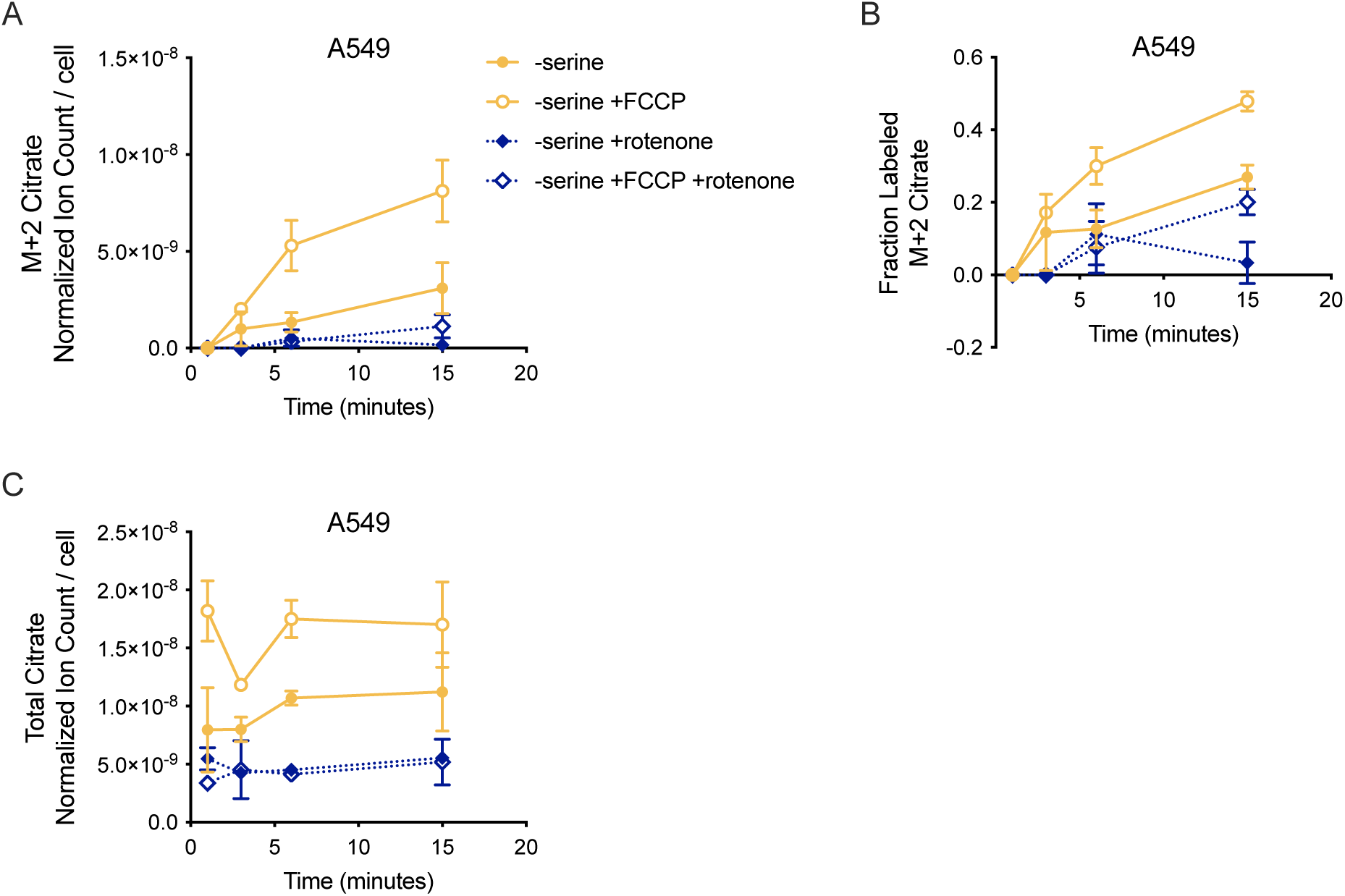
Kinetic tracing of labeled carbon from U-^13^C-glucose into citrate in cells cultured without serine and in the presence or absence of FCCP and/or rotenone. **(A)** Fraction of citrate labeled (M+2) from U-^13^C-glucose over time in A549 cells cultured without serine for 24 hours, and with or without the addition of 2 μΜ FCCP or 20 nM rotenone as indicated, n=3. **(B)** Fraction of citrate labeled (M+2) from U-^13^C-glucose over time in A549 cells cultured without serine for 24 hours, and with or without the addition of 2 μΜ FCCP or 20 nM rotenone as indicated, n=3. **(C)** Total citrate levels in A549 cells cultured without serine for 24 hours, and with or without the addition of 2 μΜ FCCP or 20 nM rotenone as indicated, n=3. All citrate measurements were normalized to internal norvaline standard and cell number. Data shown for all panels are means ± SD.

**Supplementary Figure 10.**
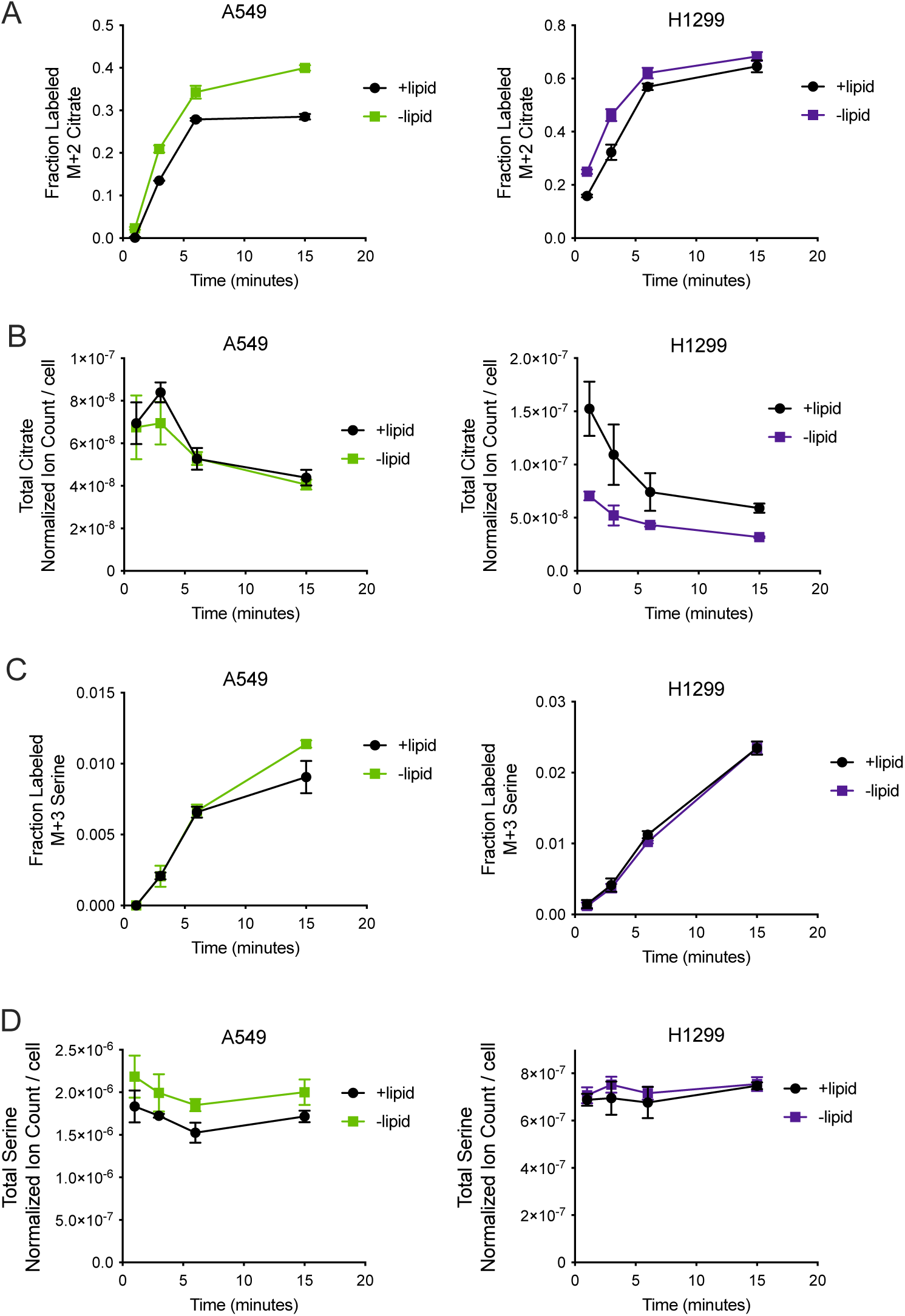
Kinetic tracing of labeled carbon from U-^13^C-glucose into citrate in cells cultured in the presence or absence of lipids. **(A)** Fraction of citrate labeled (M+2) from U-^13^C-glucose over time in A549 and H1299 cells cultured with or without lipids for 24 hours as indicated, n=3. **(B)** Total citrate levels in A549 and H1299 cells cultured with or without lipids for 24 hours for each time point shown in (A), n=3. **(C)** Fraction of serine labeled (M+3) from U-^13^C-glucose over time in A549 and H1299 cells cultured with or without lipids for 24 hours as indicated, n=3. **(D)** Total serine levels in A549 and H1299 cells cultured with or without lipids for 24 hours for each time point shown in (C), n=3. All citrate and serine measurements were normalized to internal norvaline standard and cell number. Data shown for all panels are means ± SD.

**Supplementary Figure 11.**
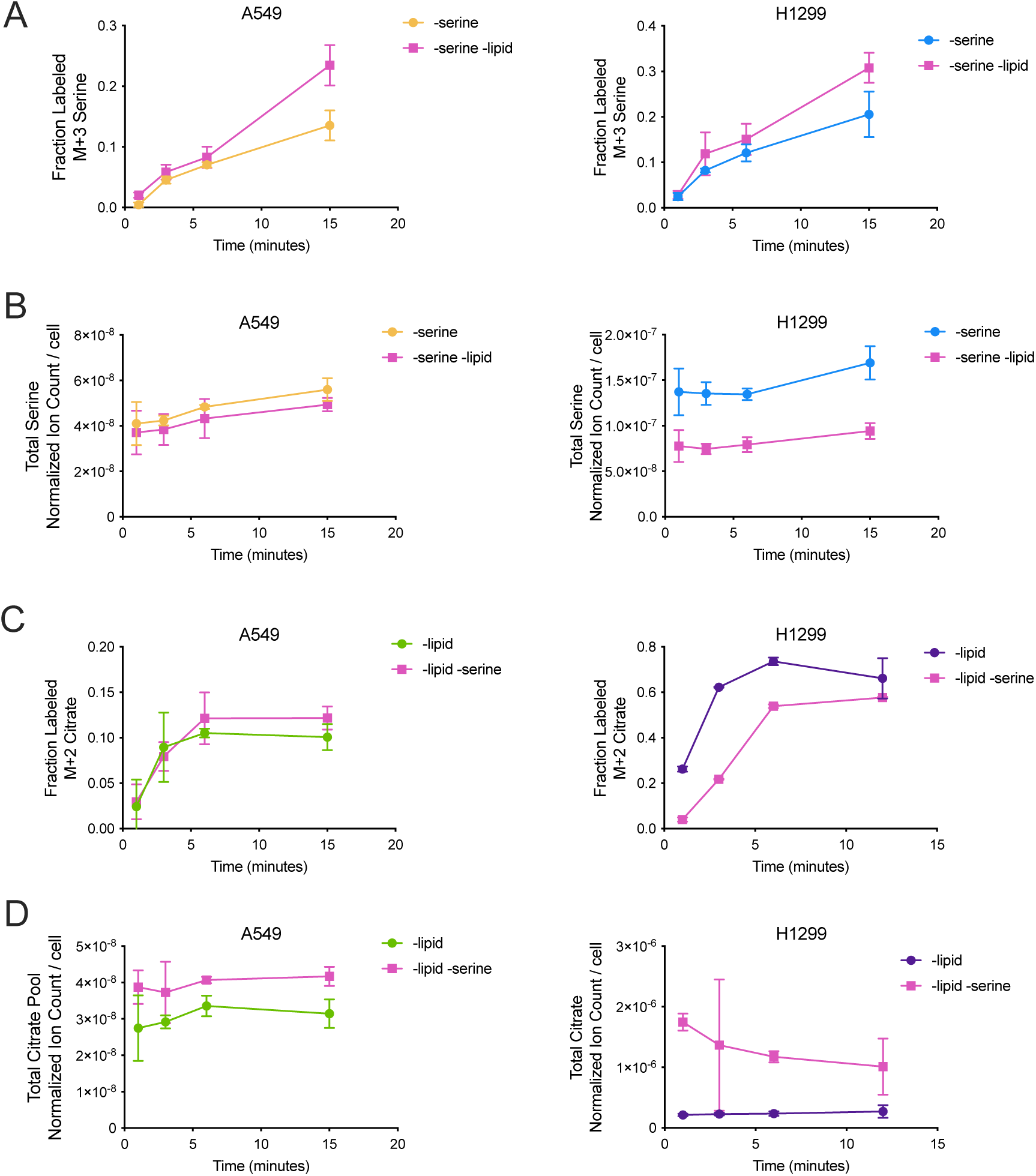
Kinetic tracing of labeled carbon from U-^13^C-glucose into serine and citrate in cells cultured in the absence of serine, lipids, or both serine and lipids. **(A)** Fraction of serine labeled (M+3) from U-^13^C-glucose over time in A549 and H1299 cells cultured without serine, and with or without lipids for 24 hours as indicated, n=3. **(B)** Total serine levels in A549 and H1299 cells cultured without serine, and with or without lipids for 24 hours for each time point shown in (A), n=3. **(C)** Fraction of citrate labeled (M+2) from U-^13^C-glucose over time in A549 and H1299 cells cultured without lipids, and with or without serine for 24 hours as indicated, n=3. (**D)** Total citrate levels in A549 and H1299 cells cultured without lipids, and with or without serine for 24 hours for each time point shown in (C), n=3. All citrate and serine measurements were normalized to internal norvaline standard and cell number. Data shown for all panels are means ± SD.

**Supplementary Table 1.**
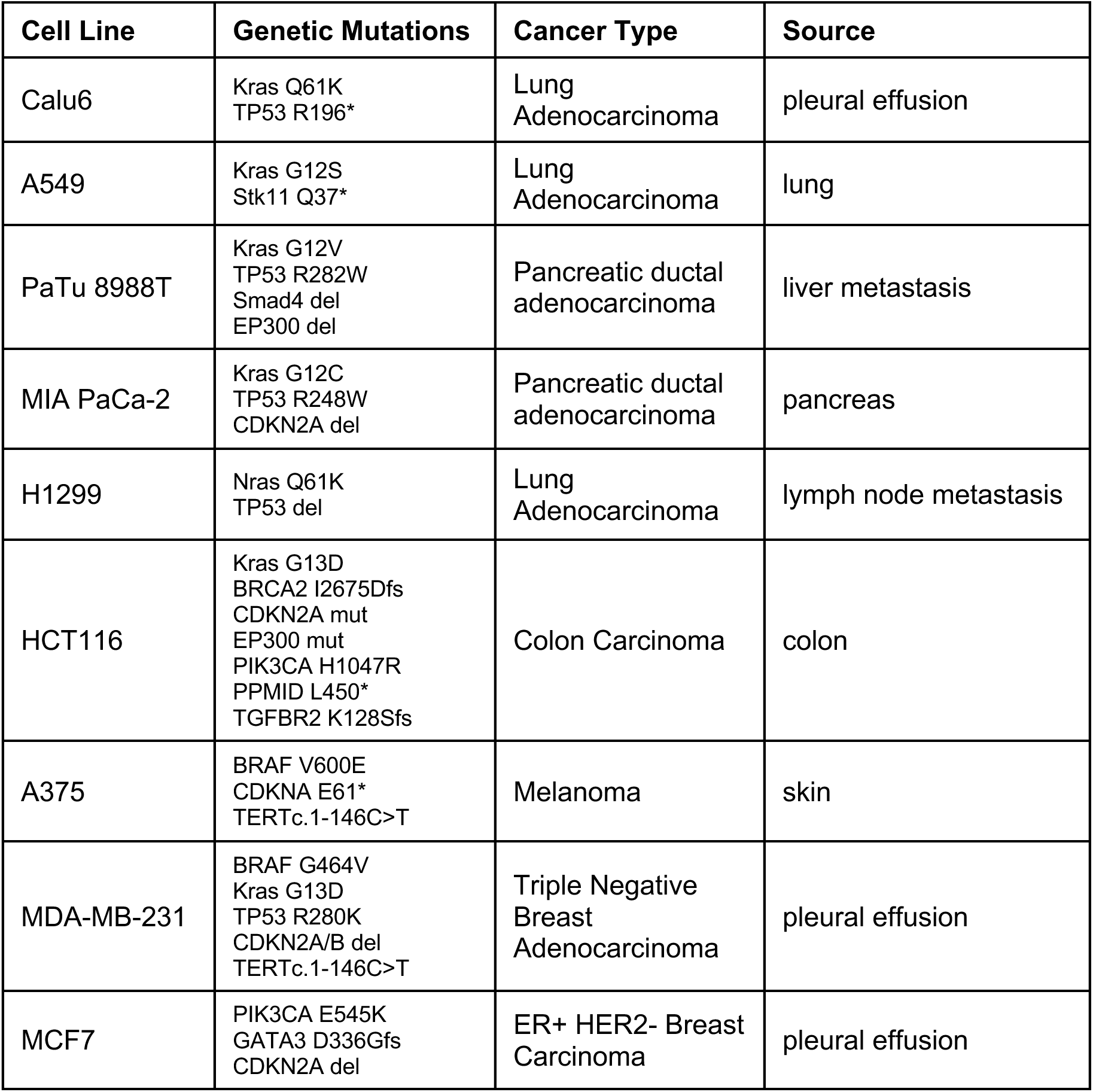

